# Neuronal silencing through depolarization block underlies odor discrimination

**DOI:** 10.64898/2025.12.11.693839

**Authors:** David Tadres, Philip Wong, Mariana Lopez-Matas, Naomi Khosalim, Moraea Phillips, Jeff Moehlis, Matthieu Louis

## Abstract

The olfactory system enables animals to distinguish between attractive and aversive odor sources through the perception of odorant molecules. A fundamental challenge is maintaining stable odor identity perception despite fluctuating stimulus intensities during navigation. Using the numerically simple olfactory system of the *Drosophila* larva, we combined a quantitative odor discrimination assay, electrophysiology, computational modelling, and ectopic expression of odorant receptors to study the encoding of odor identity. We found that depolarization block, which silences olfactory sensory neurons (OSNs) at odor concentrations 3-4 log units above the detection threshold, enables odor discrimination by creating distinct patterns of OSN activity. In a minimal system with two functional OSNs, larvae discriminate between odors when depolarization block creates differential OSN silencing but fail to distinguish odors producing qualitatively similar activity patterns in both OSNs. This principle extends to odor mixtures and the fully functional olfactory system, where the high affinity of the OR42b odorant receptor for one of the test odors enables discrimination regardless of which OSN expresses OR42b. Our results indicate that depolarization block serves as an integral feature of olfactory coding alongside combinatorial OSN activation. Selective silencing of high-affinity OSNs creates distinct neural representations and enables discrimination between odors that would otherwise produce similar activity patterns. This mechanism appears evolutionarily conserved across sensory systems, from olfactory sensory neurons to photoreceptors, suggesting that selective neuronal silencing serves as a mechanism for maintaining distinct sensory representations across stimulus intensities.

## Introduction

Animals rely on their sense of smell for essential behaviors such as locating food, identifying mates, and avoiding predators ^1^. These behaviors require navigating odor gradients by moving toward or away from an odor source based on concentration changes. A fundamental challenge in olfactory navigation is perceptual persistence: the ability to maintain a stable perception of odor identity despite fluctuating stimulus intensity. How does the peripheral olfactory system achieve this dual encoding of both odor identity and intensity?

In both vertebrates and the vinegar fly *Drosophila melanogaster*, odors are encoded through combinatorial activation of olfactory sensory neurons (OSNs) ^2–5^. Each odorant receptor (OR) is expressed in a single OSN type, and all OSNs expressing the same OR project to the same glomerulus in the antennal lobe of flies ^6,7^ or the olfactory bulb of vertebrates ^8,9^. Individual ORs can detect multiple ligands, with different ORs responding to distinct but overlapping sets of odors. A relatively small number of glomeruli can thus encode a vast array of odors, including complex natural mixtures like fruit volatiles ^3,10–17^.

The dose responses of OSNs follow an S-shaped (sigmoidal) dose-response curve that is conserved across species ^14,18–20^, a fundamental characteristic of sensory systems. These responses are typically described by Hill equations, which capture the cooperative binding dynamics of odorant-receptor interactions ^14,21^. When odor concentration exceeds the sensory threshold, OSN activity increases non-linearly and monotonically before reaching a plateau. The sensitivity of each OSN is characterized by the EC50, the concentration that elicits half-maximal response.

In olfactory systems ranging from flies to mice, the sensitivities of OSNs to a given odor follow a power law, which ensures detection across many orders of magnitude ^14,22^. This principle creates a fundamental challenge: as the concentration of an odor increases, it activates an expanding and increasingly overlapping population of OSNs with different sensitivities, potentially compromising the stability of odor perception.

The concentration-dependent recruitment of OSNs means that the representations of odor identity and intensity are intertwined in the peripheral olfactory system during navigation and olfactory tasks ^12,20,23^. This entanglement raises two central questions about combinatorial coding: Is the specific pattern of OSN activation essential for encoding odor identity? How do animals maintain stable odor recognition despite concentration-dependent changes in OSN recruitment patterns?

Unlike vision and audition, where quality perception (color/pitch) remains stable across intensity changes ^24^, olfactory perception can shift dramatically when concentrations exceed a certain range. While perceived odor identity often remains constant over moderate concentration ranges in humans, significant increases can alter both quantitative and qualitative perceptual properties ^25,26^. Similar concentration-dependent perceptual shifts including changes from attraction to avoidance behaviors, have been observed across species. In bees ^27–29^ and *Drosophila* ^30–32^, animals trained to recognize an odor at a given concentration can recognize the same odor at half or double the odor concentration. However, animals fail to recognize the same odor at much higher concentrations. What mechanisms explain this combination of concentration-invariance within certain ranges and perceptual switches beyond them?

In *Drosophila* larvae, we recently discovered that depolarization block constrains OSN activity to a range spanning approximately three orders of magnitude in concentration ^33^. When odor concentrations exceed this range, the inactivation of sodium channels silences these neurons. This phenomenon extends beyond larvae, as it has been documented in adult flies ^34,35^ and mammalian olfactory systems ^36^. Depolarization block challenges the traditional view of olfactory coding. Instead of a continuous increase in the number of active OSNs with higher concentrations, high-affinity OSNs become inactive, leading to a shift in the active OSN population. This dynamic balance between silencing and activation could serve two purposes: preserving sparse neural coding while maximizing coding capacity, and conserving energy. As some OSNs become silent, others become active, ensuring efficient use of the metabolic and computational resources of the olfactory system.

Given the essential role of high-affinity “primary” OSNs in olfactory processing ^36^, we hypothesized that their silencing through depolarization block could alter odor perception. To test this idea, we used the larval olfactory system, which offers unique advantages to study peripheral odor coding. The system consists of just 21 OSNs, each expressing a private odorant receptor (occasionally two) along with the ubiquitous co-receptor ORCO ^37–39^. Unlike most olfactory systems where multiple OSNs express the same OR, in the larval system each OR is expressed by only a single OSN. The receptive fields of larval OSNs have been extensively characterized through electrophysiology and calcium imaging ^13,14,38,40–42^. The minimal architecture of the larval olfactory system makes it ideal for studying the contribution of intrinsic neuronal mechanisms like depolarization block, while its relatively simple circuit organization enables a systematic analysis of how changes in individual OSN responses relate to odor discrimination across concentrations.

Our central hypothesis is that odor representations are shaped by both active OSNs and those that do not respond, either because they remain at basal activity due to subthreshold concentrations or because they become silent through depolarization block. This predicts that odors creating distinct patterns of non-responding OSNs should be discriminable. To test this hypothesis, we first use larvae engineered with only two functional OSNs to demonstrate that depolarization block enables discrimination between both pure odorants and odor mixtures. We then extend these findings to larvae with fully functional olfactory systems, thereby establishing that the selective silencing of specific OSNs through depolarization block can be necessary for odor discrimination.

## Results

### Depolarization block enables odor discrimination in a minimal two-OSN system

To investigate whether depolarization block enhances odor discrimination, we engineered a simplified olfactory system that retains the capacity for combinatorial coding. We genetically engineered larvae with just two functional OSNs ^42^ and searched for odor pairs that would differentially activate these neurons, with one odor inducing depolarization block in only one OSN. Based on previous work ^13,14,33^, we screened odor candidates to find pairs that would either activate both OSNs similarly (overlapping dose-response curves) or differentially silence one OSN through depolarization block (non-overlapping dose-response curves).

We generated larvae with simplified olfactory systems by rescuing expression of the odorant co-receptor ORCO in specific OSNs within an anosmic *Orco* null (*Orco^-/-^*) background ^37,39^. Using the UAS/Gal4 expression system, we drove *Orco* expression under OR-specific promoters, an approach previously validated for behavioral studies ^39,42–44^.

The *Or42b*-expressing OSN undergoes depolarization block in response to ethyl butyrate (EtB) ^33^. EtB also strongly activates the *Or42a*-expressing OSN, with signs of depolarization block at higher odor concentrations than the *Or42b* OSN ^42^. We therefore focused on the *Or42a* and *Or42b* OSNs. Previous physiological and behavioral evidence suggested ethyl acetate as another promising candidate that elicits strong activity in both the *Or42a* and *Or42b* OSNs ^13^, and the results of cross-adaptation experiments ^45^ led us to identify methyl acetate as a third candidate odor.

We characterized OSN responses using electrophysiology ^33,46^ while delivering precisely controlled odor stimuli through a microfluidic device (Figure 1A). We measured responses to 20-second stationary pulses of methyl acetate (MA), ethyl acetate (EtA), and ethyl butyrate (EtB) across several orders of magnitude in concentrations (Figure 1B-F). While all three odors activated both OSNs, their sensitivities differed markedly. The initial phasic responses showed typical sigmoidal dose-response curves (Figure 1D-F, top row). However, the steady-state responses after 15 seconds revealed inverted-U-shaped (∩) curves, where odor concentrations significantly higher than the sensory threshold silenced the OSNs through depolarization block (Figure 1D-F, bottom row).

**Figure 1:**
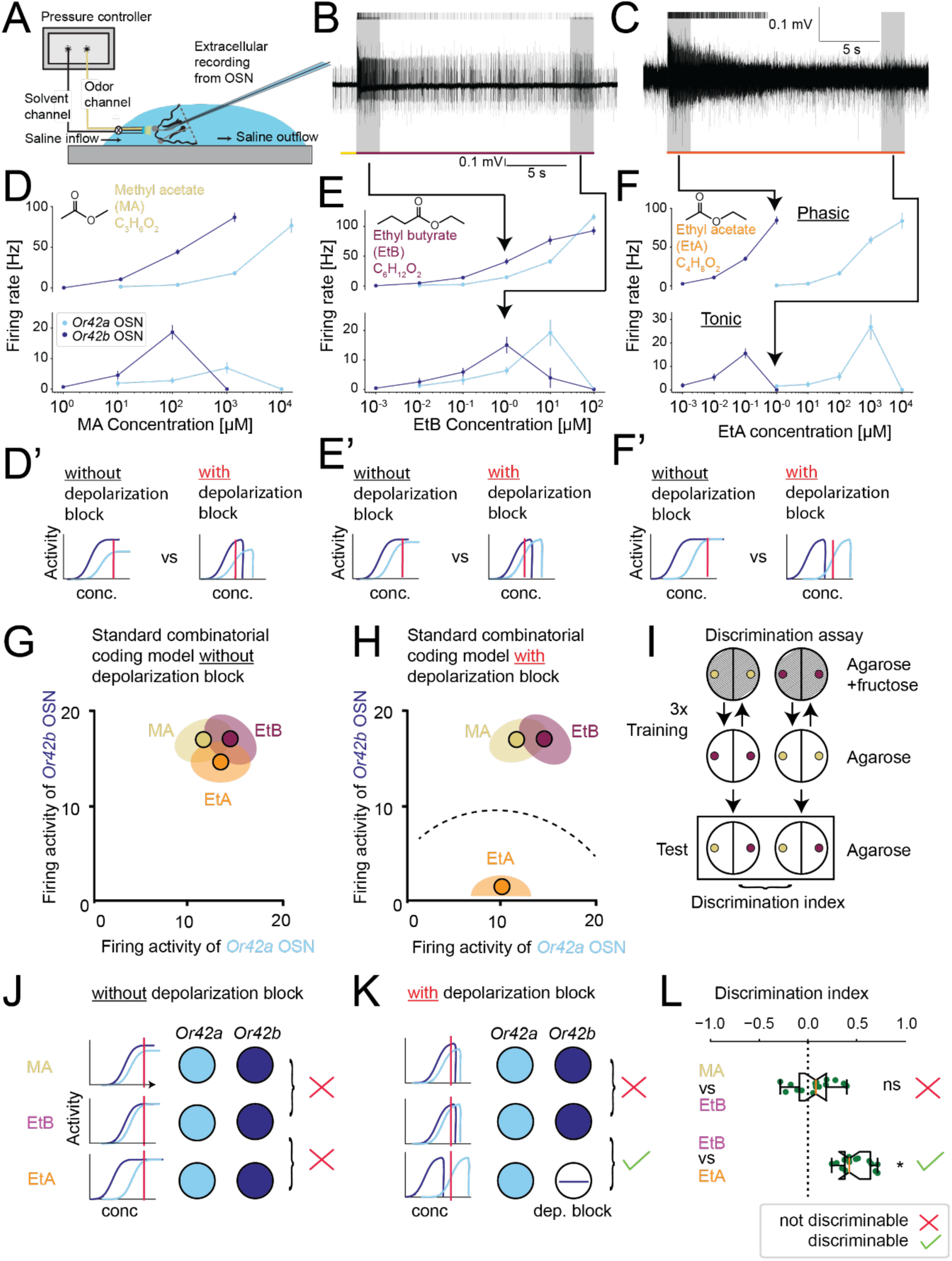
Depolarization block enhances odor discrimination capacity. **(A)** Electrophysiology setup. **(B)** Example recording of *Or42b* OSN stimulated with 1 μM EtB. Gray areas indicate timepoints for phasic (top of D/E/F) and tonic (bottom of D/E/F) dose-response curves. Vertical lines on top indicate identified spikes. Red horizontal line indicates odor stimulation (data from ref. ^33^) **(C)** Example recording of *Or42b* OSN stimulated with 1 μM EtA. Note that the *Or42b* OSN undergoes depolarization block halfway through the stimulation. **(D-F)** Phasic (top) and tonic (bottom) dose-response curves for *Or42a* OSN (cyan) and *Or42b* OSN (navy) in response to (D) MA, (E) EtB, and (F) EtA (*Or42b* OSN EtB data reproduced from ref. ^33^). **(D’-F’)** Schematic representation of the predicted neuronal activity without and with depolarization block at the concentration indicated with a red vertical bar. During prolonged EtA stimulation (F’), *Or42a* and *Or42b* OSNs cannot fire simultaneously due to the non-overlapping nature of their dose-responses. **(G)** Hypothetical firing rates without depolarization block. **(H)** Hypothetical firing rates with depolarization block. When *Or42a* OSN fires in response to EtA, *Or42b* OSN undergoes depolarization block during prolonged odor exposure. **(I)** Schematic of associative learning assay. One group experiences odor 1 with fructose reward and odor 2 without reward; a second group receives reciprocal training. Larvae discriminating between odors will navigate toward the reinforced odor during testing. **(J)** Without depolarization block, *Or42a+b* double functional (DF) larvae are predicted to fail discrimination for both MA vs. EtB and EtB vs. EtA. **(K)** With depolarization block, larvae are predicted to discriminate between EtB and EtA. **(L)** *Or42a+b* DF larvae discriminate EtB vs. EtA but not MA vs. EtB (n=15, one-sample *t* test followed by Holm-Bonferroni correction, *p<0.05). The discrimination index is defined in Methods.

Both OSNs underwent depolarization block approximately three orders of magnitude above their detection thresholds. Importantly, EtA produced non-overlapping dose-response curves: at low EtA concentrations, the *Or42b* OSN responded while the *Or42a* OSN remained at basal activity. At higher EtA concentrations, the *Or42a* OSN responded while the *Or42b* OSN entered depolarization block (Figure 1F & 1F’). By contrast, MA and EtB produced similar dose-response patterns in both OSNs from low to high concentrations (Figure 1D-E & 1D’-E’). We confirmed these physiological findings behaviorally with larvae having a single functional OSN expressing either *Or42a* or *Or42b* (Figure 1-S1A-D).

Our electrophysiological characterization suggested that non-overlapping OSN activity patterns enable odor discrimination. Given the overlapping activation patterns elicited by MA and EtB (Figure 1D versus 1E), we hypothesized that larvae with only the *Or42a* and *Or42b* OSNs functional would be unable to discriminate between these two odors. Conversely, EtB should be discriminable from EtA due to its concentration-dependent differential activation of the *Or42a* and *Or42b* OSNs (Figure 1E versus 1F). This prediction specifically depends on depolarization block (Figure 1G-H), as the sigmoidal phasic responses alone would not create such distinct activity patterns.

To test these predictions, we adapted a standard associative learning paradigm in which larvae were trained to associate odors with an appetitive sugar reward (fructose) (Figure 1I, Methods) ^47,48^. Initial experiments with larvae having a single functional OSN expressing *Or42a* or *Or42b* (hereafter *Or42a* SF or *Or42b* SF) showed that a single OSN was insufficient for odor discrimination (Figure 1-S1H). Control experiments confirmed that these animals were able to detect the tested odors and to form and retrieve memory (see Methods, Figure 1-S1E-L). These results indicated that larvae require more than one OSN type to discriminate odors.

Next, we used larvae with both OSNs functional (hereafter *Or42a+b* double functional, or *Or42a+b* DF). We compared the ability to discriminate between odors that activated both OSNs (MA and EtB) and odors that created differential OSN activity through depolarization block (EtA and EtB) (Figure 1H). As predicted, *Or42a+b* DF larvae successfully discriminated between EtA and EtB but failed to discriminate between MA and EtB (Figure 1L and Figure 1-S2E-F). Control experiments showed that each odor could be associated with reward when tested against no odor (Figure 1-S2B-D). To ascertain the contribution of OSNs to odor discrimination, we tested *Orco^-/-^* null mutant larvae and found they showed no attraction to any of the three odors at the tested concentrations (Figure 1-S2G). These results provided strong evidence that depolarization block enables odor discrimination through differential OSN activation patterns.

### Depolarization block in the *Or42b* OSN creates qualitatively distinct odor representations during discrimination

During odor discrimination tests, larvae navigate concentration gradients to reach their learned odor target (Figure 1I). To establish whether depolarization block underlies successful discrimination, we examined OSN activity patterns corresponding to actual larval trajectories toward different odor sources (Figure 2). We reconstructed the odor concentration landscape produced by two distinct sources in the test assay using a biophysical diffusion model (see Methods). We fit model parameters to concentration profiles measured through infrared spectroscopy (Figure 2-S2). Using trajectory data from real larvae, we replicated these concentration changes with a microfluidics system ^33,46^, while recording OSN activity through extracellular electrophysiology (Figure 1A).

**Figure 2:**
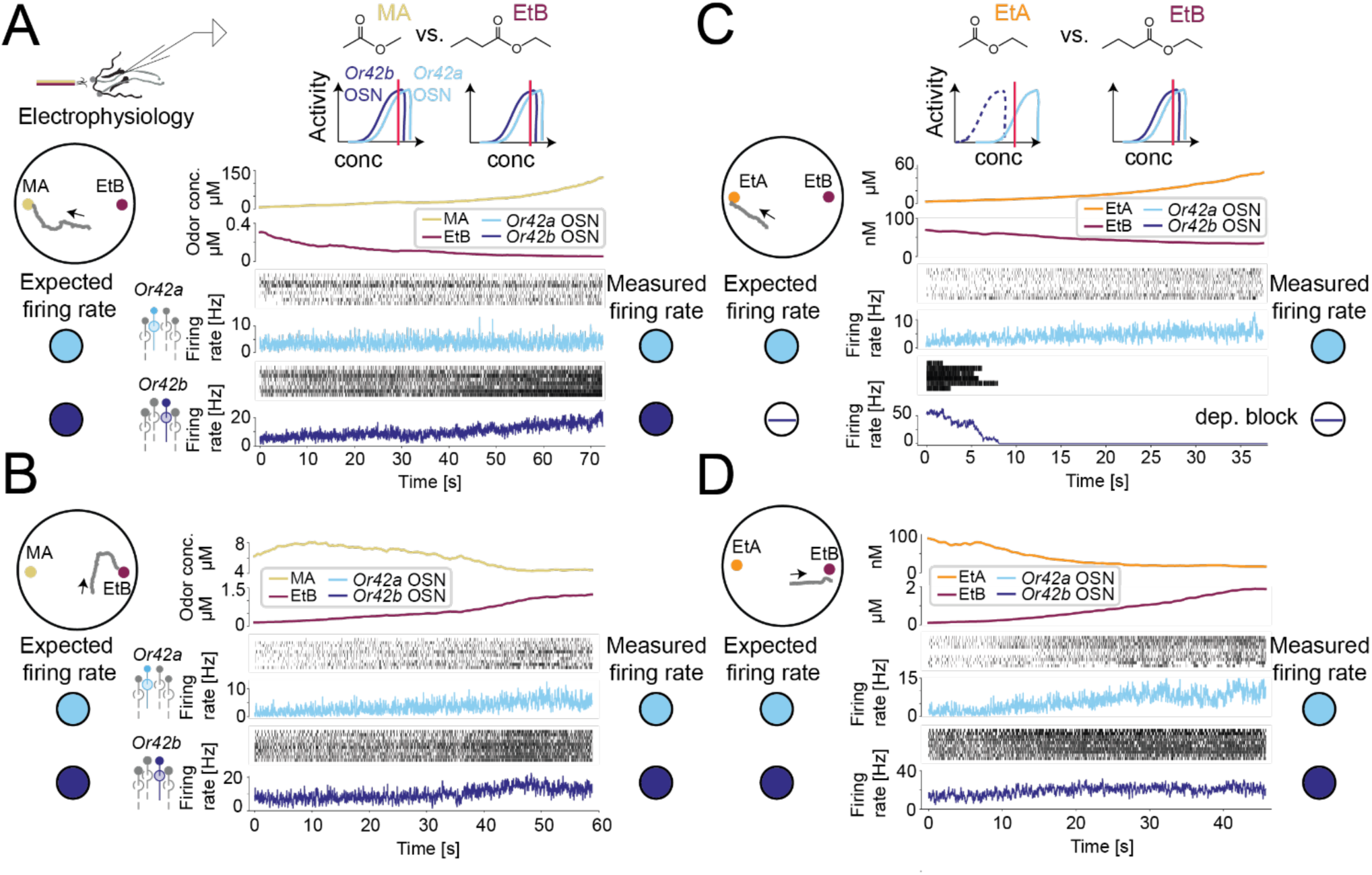
Predicted and measured OSN activity during a discrimination task. **(A)** Behavior and electrophysiology with single odors predicts *Or42a* and *Or42b* OSN activity during larval navigation toward MA or EtB (Figure 1). Replaying combined MA + EtB concentrations experienced by a representative larva moving towards MA (arrow indicates movement direction) increases firing rates in both *Or42a* OSN (n=9) and *Or42b* OSN (n=8). **(B)** Both OSNs increase firing rates when presented with MA + EtB concentrations experienced by a larva moving toward EtB (n=10 each). **(C, D)** Electrophysiology and behavior predict *Or42b* OSN enters depolarization block when larvae approach EtA (see Figures 1 and 3-S1). Replaying EtA + EtB at concentrations experienced by a larva moving toward EtA elicits activity only in *Or42a* OSN (n=10), as *Or42b* OSN (n=6) rapidly enters depolarization block. Both *Or42a* OSN and *Or42b* OSN increase firing when a larva approaches EtB (n=10 each).

When larvae approached either MA or EtB sources, both the *Or42a* and *Or42b* OSNs remained active, though with different relative activity levels (Figure 2A-B). However, when larvae approached EtA during EtA versus EtB discrimination, the *Or42b* OSN rapidly entered depolarization block (Figure 2C). When moving toward EtB instead, both OSNs increased their activity. These results support our hypothesis that larvae discriminate odors based on qualitative differences in OSN activity patterns created by depolarization block, rather than quantitative differences in relative OSN activity levels.

To analyze OSN activity patterns during learning, we applied our odor diffusion model to reconstruct odor concentrations experienced by single larvae during training (Figure 2-S1). We adapted a biophysical model ^33^ to predict OSN responses to these dynamic concentration changes (Figure 3). The model parameters were optimized to fit both static stimulation pulses with and without depolarization block (Figures 3-S2 to 3-S4), and dynamic concentration changes relevant to navigation (Figure 3-S5), while maintaining the same parameters for the spike generation module across all conditions. The model accurately predicted OSN responses to trajectory-based stimuli (Figure 3B-C).

**Figure 3:**
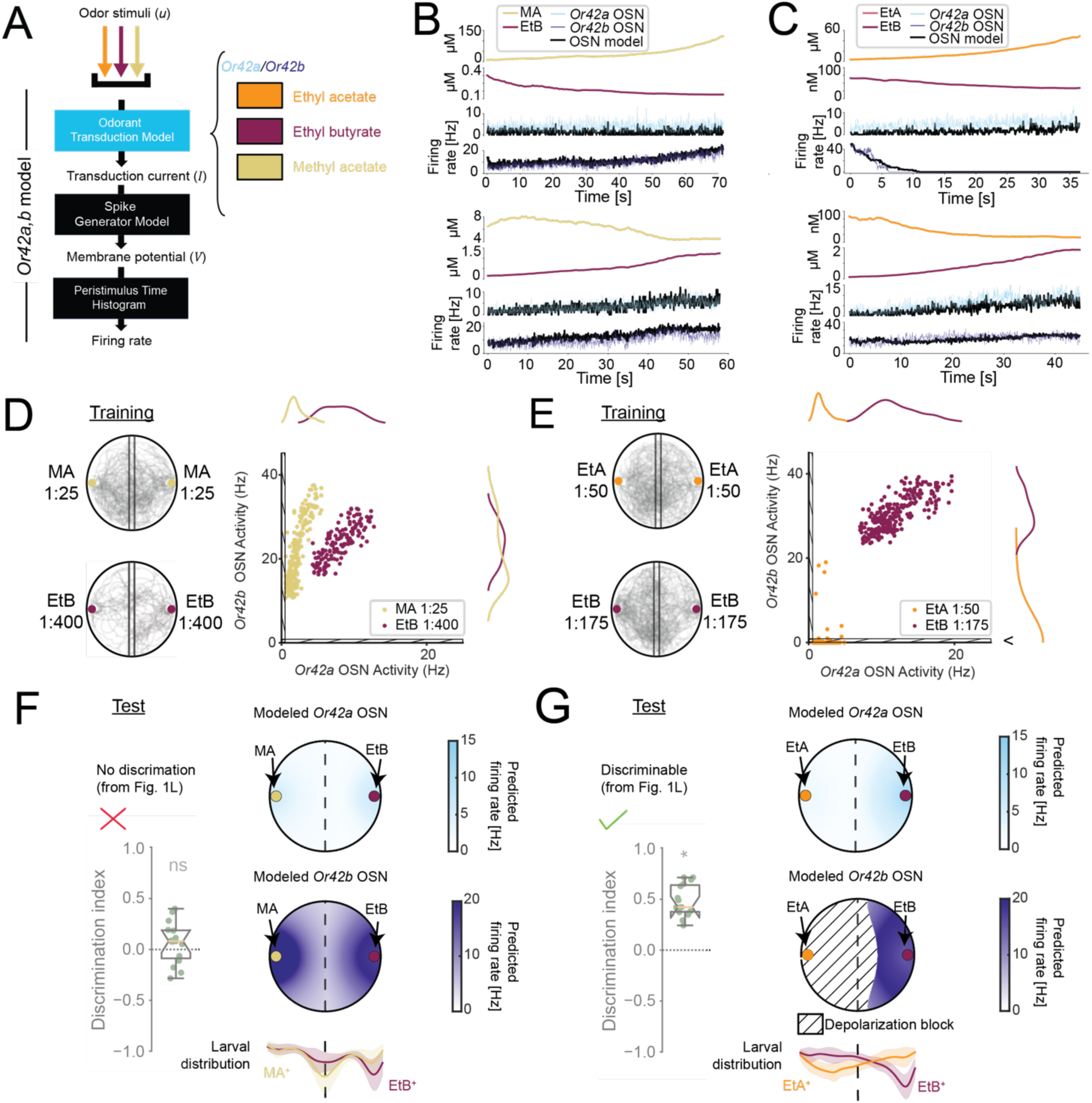
OSN model predictions show depolarization block enables discrimination. **(A)** Framework of the integrated OSN model ^33^ with an adapted odorant transduction module accounting for *Or42a* and *Or42b* OSN differences and odor-specific responses. See Figures 3-S2 to 3-S5 for model fitting data. **(B, C)** OSN model predictions during discrimination (black lines, see Figure 2). **(D)** Left: Trajectory segments during the last training session associating MA (top) and EtB (bottom) with reward. Right: Simulated final OSN activity during training. Each point represents the average activity of *Or42a* and *Or42b* OSN of a single larva during the last 5 seconds of the training (MA 1:25 vs. EtB 1:400: n=204 and n=150 larvae, respectively; EtA 1:50 vs. EtB 1:175: n=231 and n=243, respectively). Both *Or42a* and *Or42b* OSN remain active during training with MA and EtB. **(E)** Left: Trajectory segments during training with EtA (top) or EtB (bottom). Right: Simulated final OSN activity. For most larvae trained with EtA, *Or42b* OSN undergoes depolarization block (hatched region indicates sub-baseline firing <1.3Hz). **(F)** Left: Discrimination index showing MA and EtB are not discriminable (from Figure 1L). Top: Predicted OSN activity during test shows both *Or42a* (light blue) and *Or42b* (dark blue) OSNs active throughout the dish. Bottom: Larval spatial distributions during test (solid line: mean; shaded area: SD; yellow: MA+ training; red: EtB+ pairing). **(G)** Left: Discrimination index showing EtA and EtB are discriminable (from Figure 1L). Top: Predicted OSN activity shows *Or42b* OSN in depolarization block across more than half of the dish. Bottom: Larvae trained with EtA (EtA^+^, light red) accumulate where *Or42b* OSN undergoes depolarization block. Electrophysiology sample sizes given in Figure 1.

During training with MA and EtB, both OSNs maintained above-baseline activity with quantitative differences in their relative activation levels (Figure 3D). During testing, larvae showed similar spatial distributions regardless of whether they were trained to approach MA or EtB (Figure 3F), consistent with their inability to discriminate these odors (Figure 1L). By contrast, training with EtA versus EtB produced categorically distinct OSN activity patterns: EtB activated both OSNs while EtA activated the *Or42a* OSN and induced depolarization block in the *Or42b* OSN (Figure 3E). During testing, the *Or42b* OSN remained in depolarization block near the EtA source but was strongly activated near the EtB source (Figure 3G, top and center). Larvae reliably accumulated near their trained odor source (Figure 3G, bottom), demonstrating successful discrimination. These results suggest that depolarization block enables odor discrimination by creating qualitatively distinct OSN activity patterns.

### Depolarization block shapes the perception of odor mixtures

To further test whether depolarization block changes odor perception, we examined mixtures of two monomolecular odors that each induce depolarization block in different OSNs. Within the concentration range tested, ethyl acetate (EtA) produces depolarization block specifically in the *Or42b* OSN (Figure 1F), while 4-hexen-3-one (4H3O) induces depolarization block in the *Or42a* OSN ^33^. We confirmed that 4H3O preferentially activates the *Or42a* OSN at low concentrations while activating the *Or42b* OSN only at higher concentrations (Figure 4C and Figure 4-S1). This differential sensitivity was confirmed through behavioral experiments (Figure 4-S2A).

**Figure 4:**
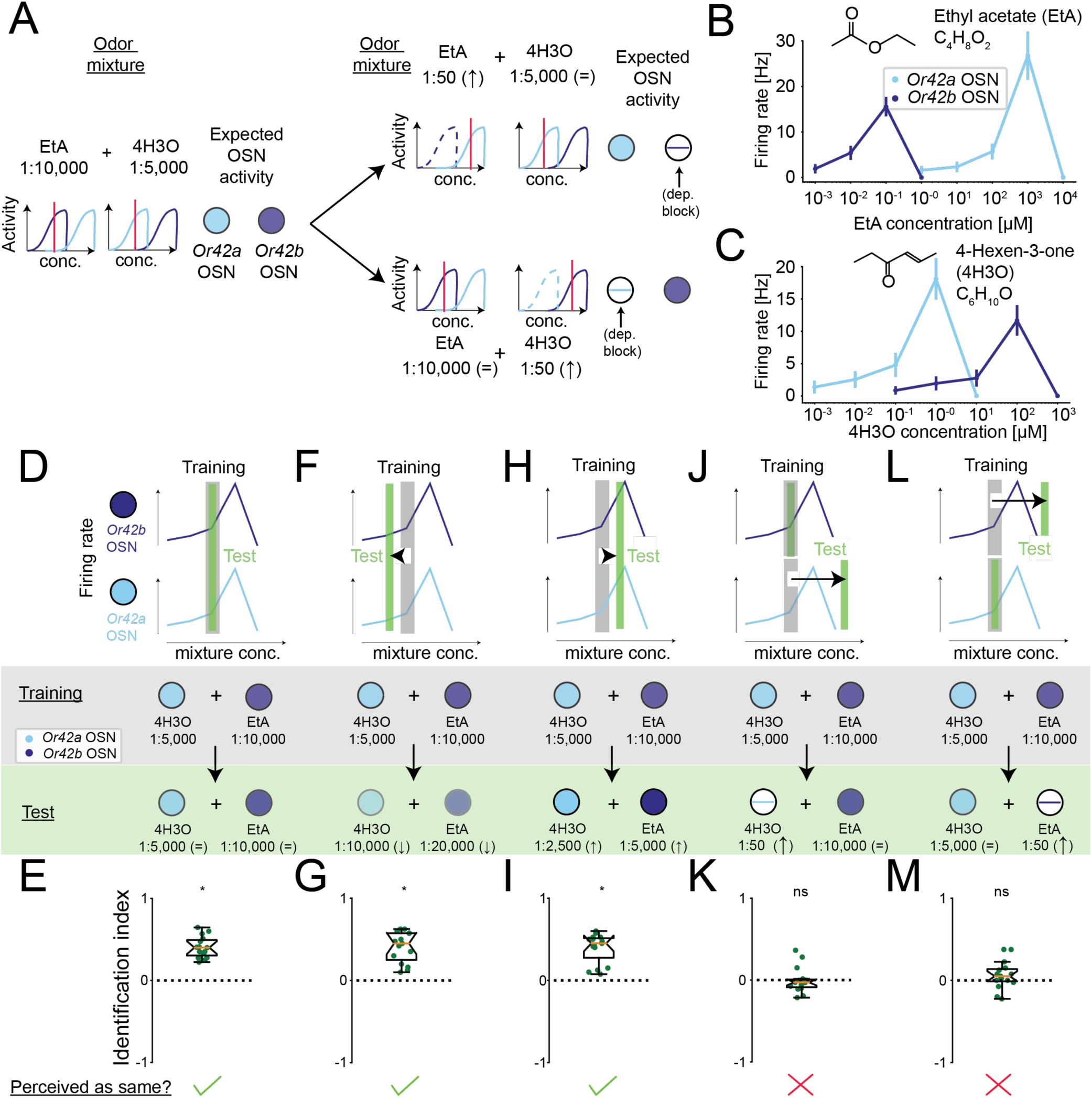
Depolarization block alters odor mixture perception. **(A)** Experimental design using odor mixtures that selectively induce depolarization block in *Or42a* or *Or42b* OSN by varying component concentrations. Red lines indicate approximate neuronal activity at each concentration; dashed lines indicate predicted depolarization block. **(B)** *Or42a* and *Or42b* OSNs show non-overlapping EtA dose-response curves (from Figure 1F). **(C)** 4-Hexen-3-one (4H3O) induces depolarization block in *Or42a* OSN at concentrations two orders of magnitude lower than Or42b OSN. **(D)** Schematic showing expected OSN activity ranges for EtA (top) and 4H3O (bottom) in *Or42b* (navy) and *Or42a* (cyan) OSNs during training (gray bar) and test (green bar). Circles indicate approximate firing rates. Training and test used identical concentrations. **(E)** *Or42a+b* DF larvae recognize a 4H3O + EtA mixture at the same concentration (n=15, one-sample *t* test). **(F)** Halving 4H3O and EtA concentrations decreases firing in both OSNs (lighter shading). **(G)** Larvae still recognize the mixture as the same (n=15, one-sample *t* test). **(H)** Doubling both concentrations increases firing in both OSNs. **(I)** Larvae recognize the mixture as identical (n=15, Wilcoxon signed-rank test). **(J)** Substantial increase in 4H3O concentration during test is predicted to induce *Or42a* OSN depolarization block. **(K)** Larvae fail to recognize the mixture (n=15, Wilcoxon signed-rank test). **(L)** Substantial increase in EtA concentration during test is predicted to induce *Or42b* OSN depolarization block. **(M)** Larvae fail to recognize the mixture (n=15, one-sample *t* test). Throughout the figure, *p<0.05. The identification index is defined in Methods.

Associative learning experiments revealed that *Or42a+b* DF larvae could recognize a mixture of EtA and 4H3O that activated both OSNs (Figure 4D; see Figure 1F for EtA and Figure 4C for 4H3O). The mixture elicited robust associative learning (Figure 4E): animals showed enhanced attraction when the mixture was paired with sugar reward and reduced attraction when paired with no reward (Figure 4-S2C). The learned response persisted when larvae were trained with the original concentration but tested with either half (Figure 4F-G and Figure 4-S2D) or double (Figure 4H-I and Figure 4-S2E) the concentration of the original mixture. Larvae can thus generalize across conditions that alter the relative levels of OSN activity while maintaining activation of both neurons.

When the concentration of either 4H3O (Figure 4J) or EtA (Figure 4L) was increased sufficiently to induce depolarization block in the *Or42a* or *Or42b* OSN, larvae failed to recognize the original mixture (Figure 4K & 4M and Figure 4-S2F-G). Schematic diagrams of OSN activity states (Figure 4D-L, middle row) summarize how depolarization block creates distinct neural representations: training with a mixture that silences one OSN prevents generalization to conditions where both OSNs are active. Importantly, all mixture conditions used during training induced comparable levels of innate attraction (Figure 4-S2B), ruling out the possibility that failed recognition resulted from reduced odor attractiveness when one OSN entered depolarization block. These results provide strong evidence that the selective silencing of OSNs through depolarization block changes the perceived odor identity of mixtures by creating qualitatively distinct activity patterns.

### Function of the OR42b receptor is essential for odor discrimination in the intact olfactory system

While the above results demonstrate the role of depolarization block in a minimal two-OSN system, *D. melanogaster* larvae possess a total of 21 OSNs that enable odor representations through combinatorial coding ^13,49^. To examine whether depolarization block remains functionally significant in the complete olfactory system, we built on our findings of EtA versus EtB discrimination in the *Or42a* and *Or42b* OSNs (Figures 1-3) and inspected the condition underlying discrimination between these odors in the fully functional olfactory system (Figure 5).

**Figure 5:**
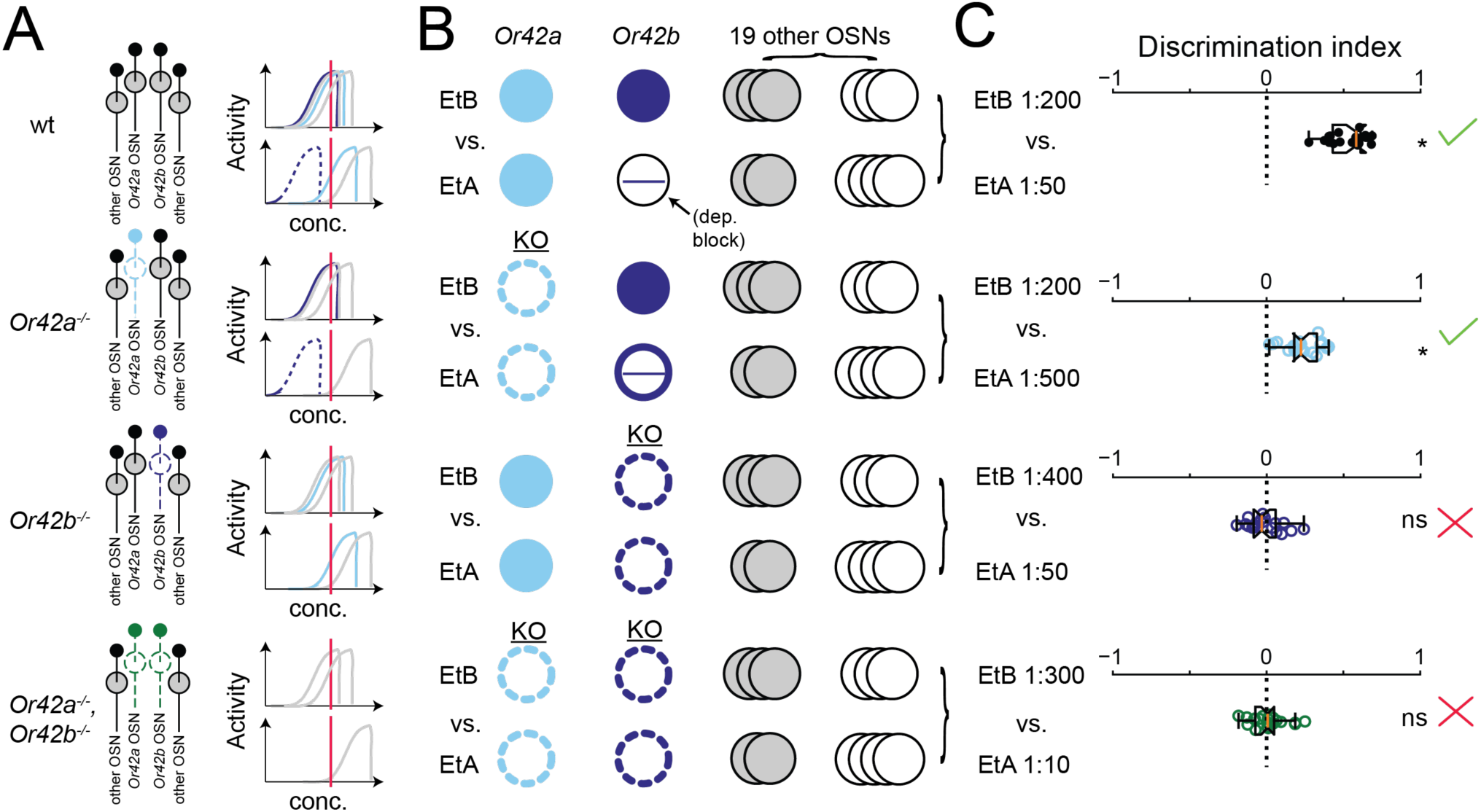
*Or42b* OSN is essential for discrimination in the intact olfactory system. **(A)** Left: Genotypes. Right: Red lines indicate approximate neuronal activity at each concentration; dashed lines indicate predicted depolarization block. Other OSN activity inferred from ref. ^14^ (Figure 5-S2). **(B)** Schematic of OSN activity patterns. **(C)** Larvae lacking OR42b receptor fail to discriminate EtA from EtB (n=20, one-sample *t* test followed by Holm-Bonferroni correction, *p<0.05). *Or42b* OSN is the most EtA-sensitive OSN and undergoes depolarization block at lower EtA concentrations than other OSN (see Figure 5-S1 and ref. ^14^).

Published calcium imaging and electrophysiology data of all 21 larval OSNs revealed that high concentrations of EtB activate multiple OSNs besides *Or42a* and *Or42b*, including *Or35a* and *Or47a* ^13,14,42^. By contrast, EtA primarily activates the *Or42b* and *Or42a* OSNs, with recruitment of other OSNs including the *Or47a* OSN only at high concentrations compared to the sensory threshold of most sensitive *Or42b* OSN ^13,14^. This activation of distinct sets of OSNs (Figure 5-S1) provides an opportunity to test the importance of depolarization block in the fully functional system.

Wild-type (*wt*) larvae successfully discriminated between EtB and EtA (Figure 5C, top row). To test the specific role of depolarization block, we examined receptor knockout mutants under conditions where both odors induced equal attraction levels to minimize discrimination based on intensity differences (Figure 5-S2A). *Or42a^-/-^* larvae remained capable of discrimination (Figure 5C, second row from top), demonstrating that the *Or42a* OSN is not required for EtA versus EtB discrimination. This result is consistent with our finding that discrimination depends on selective depolarization block of the *Or42b* OSN (Figures 2C-D and 3E): while the *Or42a* OSN responds to both odors without undergoing depolarization block, only the *Or42b* OSN undergoes differential silencing by entering depolarization block in response to EtA but remaining active for EtB. Confirming the essential role of *Or42b* OSN, *Or42b^-/-^* larvae failed to discriminate between the odors (Figure 5C, third row from top), despite the differential activation of multiple OSNs by EtB (Figure 5C and Figure 5-S2B).

We confirmed that *Or42a* and *Or42b* OSNs are essential for discriminating between EtA and EtB using larvae bearing a double knockout *Or42a^-/-^*,*Or42b^-/-^* (hereafter Del(*Or42*a,*Or42b*)). Del(*Or42*,*Or42b*) larvae failed to discriminate between EtB and EtA while maintaining equal attraction to both odors (Figure 5C, bottom row). To preserve chemotactic behavior in these mutants, it was necessary to use higher EtA concentrations to recruit OSNs that are less sensitive to EtA such as *Or47a* OSN (Figure 5-S1C) ^13,14^ to achieve equal naïve preference (methods). These results suggest that even in the fully functional olfactory system, an OSN undergoing depolarization block can be essential for odor discrimination.

### Depolarization block induced by the OR42b receptor enables discrimination in any OSN

Can any OSN contribute to odor discrimination through depolarization block when expressing an appropriate receptor, or is this property restricted to specific OSNs within the olfactory circuit? To approach this question, we ectopically expressed odorant receptors in selected OSNs of Del(*Or42a*,*Or42b*) double-mutant larvae. Odorant receptors maintain their molecular properties when expressed in non-native OSNs, a principle established through systematic characterization of OR function in the “empty” *Or22a* neurons of adult flies ^3,50,51^. We previously showed that expressing the OR42b receptor in the *Or1a* OSN is sufficient to produce EtA-induced depolarization block ^33^.

Using UAS-OR transgenes, we expressed the OR42a receptor in either the *Or42a* or *Or42b* OSN of Del(*Or42a*,*Or42b*) mutants. Neither manipulation restored discrimination ability (Figure 6A-C, top two rows). However, expressing OR42b in either the *Or42a* or *Or42b* OSN fully rescued discrimination (Figure 6A-C, bottom two rows). The restoration of discrimination through OR42b expression in the *Or42a* OSN indicates that the molecular properties of OR42b enable discrimination through depolarization block, rather than the specific identity of the OSN in which it is expressed. These results demonstrate that OR42b-mediated depolarization block provides a mechanism for odor discrimination through the creation of qualitatively distinct OSN activity patterns.

**Figure 6:**
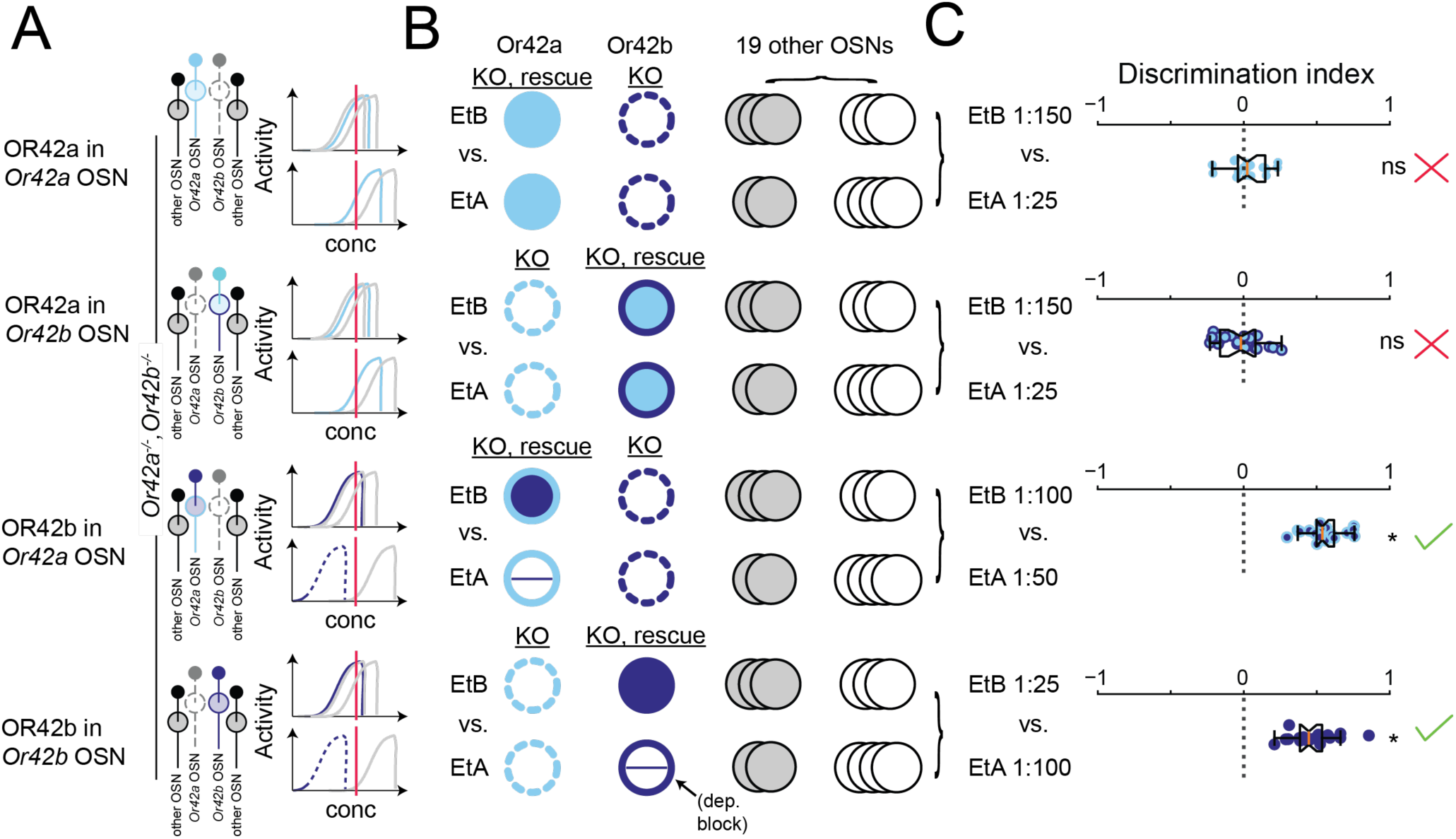
OR42b-mediated depolarization block is essential for discrimination. **(A)** Left: Genotype (all in Del(*Or42a,Or42b*) double-knockout background). Right: Red lines indicate approximate neuronal activity at each concentration; dashed lines indicate predicted depolarization block. Other OSN activity inferred from ref. ^14^ (Figure 5-S1). **(B)** Schematic of OSN activity patterns. **(C)** OR42a alone fails to rescue EtA vs. EtB discrimination in either *Or42a* (top) or *Or42b* (second from top) OSNs. Conversely, OR42b expression in either *Or42a* (third from top) or *Or42b* (bottom) OSNs rescues discrimination (n=20, one-sample *t* test followed by Holm-Bonferroni correction, *p<0.05).

## Discussion

Animals face a fundamental challenge in olfaction: maintaining stable perception of odor identity while using intensity changes to navigate toward odor sources. The olfactory coding hypothesis posits that odor percepts arise from unique patterns of neural activity in the peripheral olfactory system ^2^. This principle has been documented across species from flies to mice, where increasing odor concentrations recruit expanding populations of olfactory sensory neurons (OSNs) with different sensitivities ^3,20^. Intriguingly, early studies in mammals provided evidence that some OSNs “drop out” of odor representations at high concentrations ^12^, suggesting that odor coding involves both recruitment and selective silencing of sensory neurons. To date, the mechanistic basis and functional significance of this phenomenon has remained unclear, due in part to technical difficulties in simultaneously manipulating multiple sensory neurons and the potential role of temporal dynamics in concentration-invariant coding ^52,53^.

In *Drosophila*, OSN responses to brief odor pulses follow S-shaped (sigmoidal) dose-response curves ^13,35,41,54^. However, natural olfactory behaviors often involve sustained exposure to odors that vary both in intensity and duration. From larvae navigating toward food sources to mice following scent trails, animals must process odor information across timescales ranging from milliseconds during sniffing to seconds during source localization ^55–57^. Recent work in larvae and mice revealed that high-affinity OSNs enter depolarization block at higher concentrations during sustained odor exposure ^33,36^. This intrinsic neuronal mechanism provides an explanation for the selective silencing of OSNs observed in earlier studies and appears particularly important when animals experience prolonged exposure to high odor concentrations, such as when making decisions after reaching an odor source.

The distribution of sensitivities across the population of OSNs responding to a given odor follows a power-law in *Drosophila* ^14,22^, ensuring detection across many orders of magnitude. Depolarization block introduces a complementary mechanism for maintaining sparse representations. Instead of having an ever-expanding population of active OSNs with increasing concentrations, depolarization block provides an efficient, cell-autonomous mechanism where high-affinity OSNs become silent just as low-affinity OSNs begin responding ^33^. Depolarization block thus reveals a new dimension of odor coding, where both activation and selective silencing of OSNs shape the neural representation. The cell-autonomous nature of this mechanism ensures rapid and reliable control of OSN activity without requiring complex circuit interactions, while changes in odor concentration are further encoded by the temporal dynamics of OSN responses ^46,58,59^, contributing to a multi-dimensional representation of both odor identity and intensity.

Using a genetically engineered system with just two functional OSNs, we demonstrated that larval discrimination depends on depolarization block creating distinct activity patterns as animals navigate toward odor sources (Figure 2). When both OSNs remain active throughout the discrimination task, even substantial differences in their relative firing rates are insufficient to enable discrimination (Figure 3D). In contrast, selective silencing of specific OSNs through depolarization block enables larvae to discriminate between odors (Figure 3E). This principle extends to more complex, naturalistic conditions: when larvae encounter odor mixtures, depolarization block creates distinct neural representations that enable discrimination (Figure 4). Such qualitative changes in OSN activity patterns provide a robust coding mechanism that maintains stable odor recognition despite the variable concentrations encountered during natural behavior.

Concentration-dependent perceptual changes have been extensively documented across species ^26–32,36,60^. In insects, both bees and adult flies can recognize odors when concentrations vary moderately from the training level but fail at concentrations far above it, often in the range where depolarization block would be expected to occur (3-4 log units above detection threshold). Similar perceptual transitions occur in mammals: mice show altered behavioral responses to high odor concentrations ^36^, and humans report qualitative changes in odor perception when concentrations increase substantially ^26^. Our findings suggest that depolarization block provides a mechanistic basis for these perceptual transitions. First, when varying mixture components such that high-affinity OSNs enter depolarization block, larvae cannot recognize the original mixture (Figure 4). Second, in the intact olfactory system with 21 functional OSNs, the high-affinity receptor OR42b enables discrimination through selective silencing, independent of which OSN expresses it (Figures 5 and 6). Selective silencing of OSNs creates distinct neural representations that explain concentration-dependent perceptual changes observed across species.

Our results provide a mechanistic explanation for previously unexplained observations in *Drosophila*. For instance, odors that strongly activate specific receptors at low concentrations often trigger concentration-dependent perceptual changes ^31,32^. By contrast, odors like benzaldehyde that primarily activate low-affinity receptors ^3,13,14^, produce more stable percepts across concentrations. Depolarization block might also explain a puzzling finding in adult flies with only a single functional OSN type: their ability to discriminate between certain odor pairs ^61^. This discrimination capability could arise when one odor induces depolarization block while the other odor maintains OSN activity, creating qualitatively distinct neural states. Such a mechanism is particularly plausible given that the discriminable odor pairs in these experiments often differ in their receptor binding affinities ^3,62^.

The peripheral encoding of odors involves multiple parallel mechanisms that generate distinct neural representations ^4^. Previous studies have emphasized three key mechanisms: the combinatorial activation of OSNs ^2^, complex firing patterns in individual neurons ^52,63^, and circuit-level processing through lateral inhibition and second-order neurons in the olfactory bulb and antennal lobe ^64–67^. Single OSNs also encode information through their temporal dynamics, capturing changes in odor intensity to guide navigation ^46^. By selectively silencing high-affinity (“primary”) OSNs, depolarization block creates qualitatively distinct neural representations, enabling odor discrimination independent of circuit-level computation.

According to the primacy coding model, the most sensitive OSNs dominate initial odor representation through their order of activation ^52^. Our findings suggest that depolarization block provides a complementary mechanism operating over longer timescales. While primacy coding captures essential features of early odor processing, particularly during brief odor sampling, depolarization block enables qualitatively distinct representations during sustained exposure through selective silencing of high-affinity OSNs. The interaction between these temporal scales creates a rich coding space where both activation sequence and selective silencing shape odor perception.

The functional significance of depolarization block extends beyond the olfactory system. In the mammalian retina, intrinsically photosensitive retinal ganglion cells (ipRGCs) create distributed intensity codes through depolarization block and maintain contrast sensitivity across a wide range of light intensities ^68^. Individual ipRGCs become silent at distinct light intensities, which parallels how different OSNs enter depolarization block at specific odor concentrations. A distinct population of retinal ganglion cells relies on depolarization block to signal negative contrast, demonstrating its role in active sensory computation ^69^. More recently, neurons in the paraventricular thalamus were shown to use depolarization block during motivated behaviors, which reveals its importance in higher-order processing ^70^. The emergence of depolarization block as a coding mechanism across diverse neural circuits suggests that it represents a fundamental strategy for neural computation.

Our findings establish depolarization block as an integral feature of sensory coding rather than a pathological state. This coding mechanism operates through a cell-autonomous process that creates distinct activity patterns without requiring complex circuit interactions. It ensures reliable responses across broad stimulus intensity ranges while maintaining computational efficiency. Depolarization block appears particularly valuable in sensory systems that process information varying in both intensity and duration, from olfactory neurons tracking odor gradients to retinal cells adapting to changing light levels. The independent evolution of this mechanism across sensory modalities and brain regions suggests that selective neuronal silencing represents an efficient solution to a fundamental computational challenge: maintaining distinct neural representations despite dramatic variations in input intensity. Our work shows that depolarization block enables odor discrimination through distinct peripheral representations and reveals a role for intrinsic neuronal properties in odor perception.

## Acknowledgment

ML thanks the Vosshall lab where the Del(*Or42a*,*Or42b*) double mutant was generated. We are grateful for extensive discussions with Bertram Gerber about the larval learning assay and its adaptation for the present study. We thank David Jarriault and Elena Knoche for support with the generation of pilot data during an early phase of the project. We are grateful to Vivek Jayaram and Parvez Ahammad for analysis and modeling of the spike trains produced by an initial set of recordings of the *Or42a* and *Or42b* OSNs. We thank Tim Currier for discussion and comments on the manuscript.

## Methods

### Fly stocks

Larvae were grown at 22°C and 60% humidity in 12-hour light/12-hour dark conditions. Throughout the text, *D. melanogaster* “wild type” refers to w^1118^. Flies with the UAS-ChrimsonR transgene ^71^ were raised on fly food containing 0.5 mM all-*trans*-retinal (Sigma-Aldrich, R2500) protected from light.

**Table S1:**
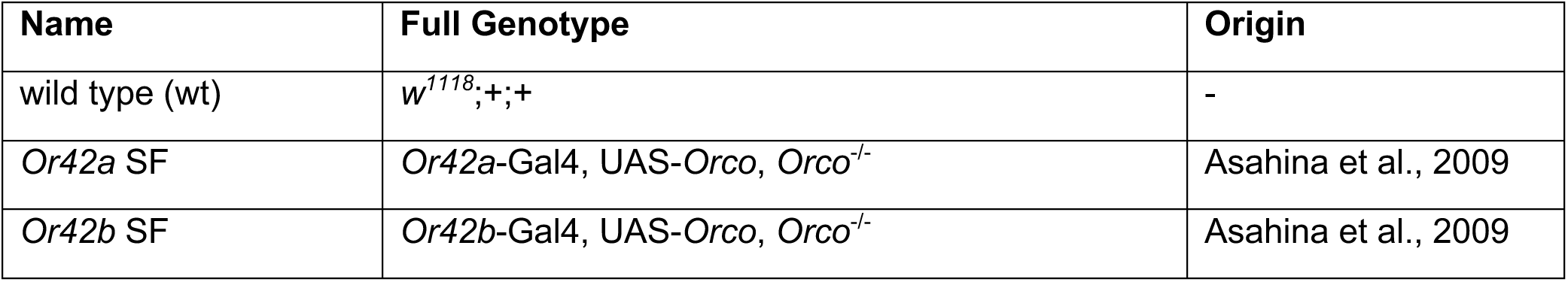
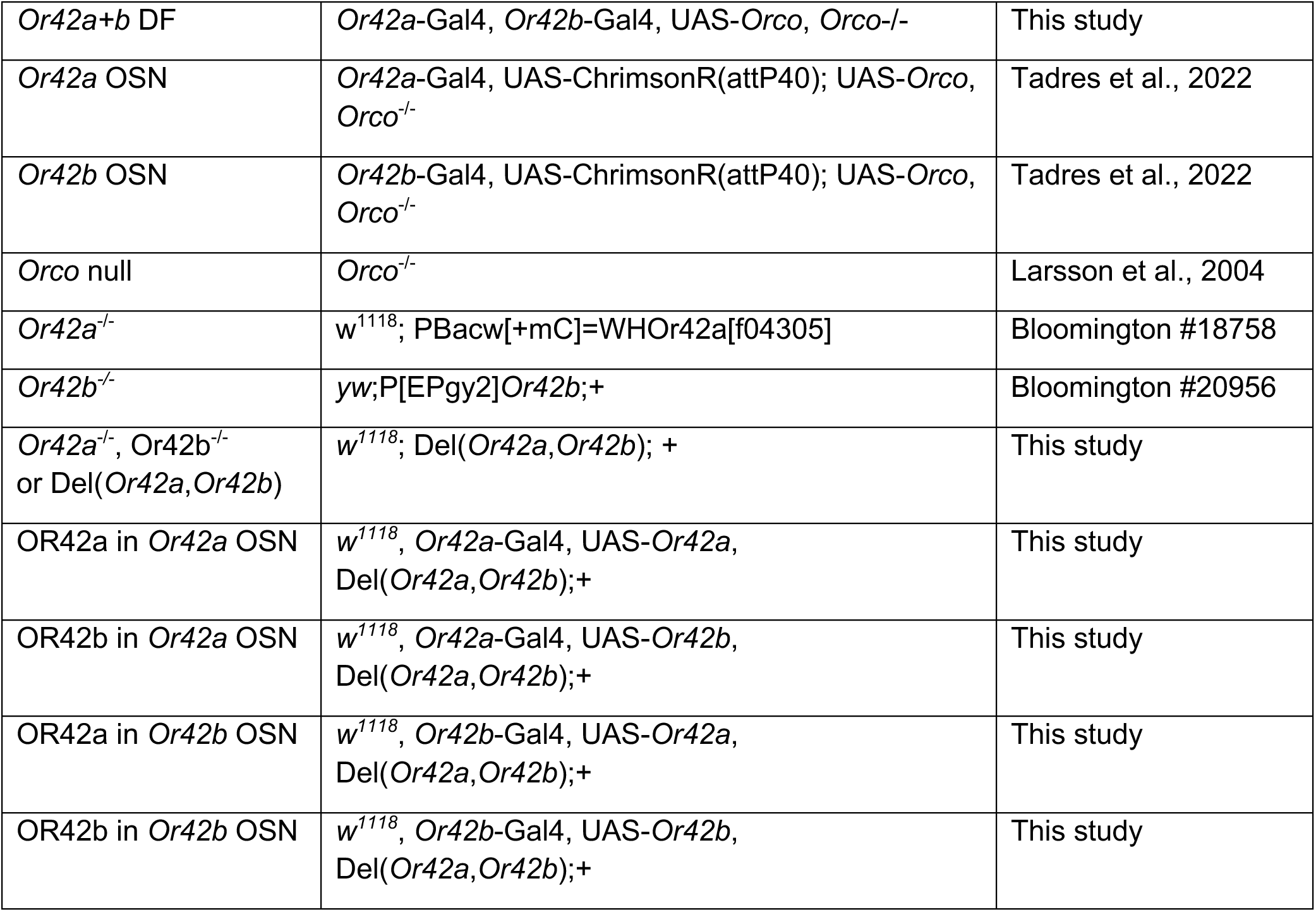
List of genotypes used in this study.

### Reagents

Only odors of the highest purity were used:

**Table S2:**
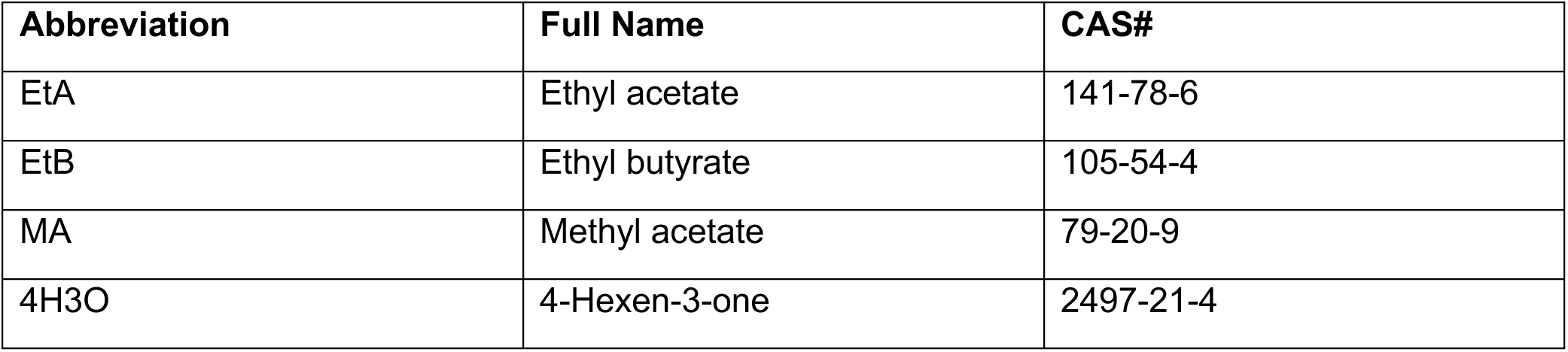
List of odors used in this study.

### Electrophysiology

Unless indicated otherwise, electrophysiological recordings and odor delivery was performed according to a protocol described in refs. ^33,46^. Each animal was used for several stimulation sets (for example, step, linear ramp and exponential stimuli) with a maximum of two recordings per stimulus type (for example, two step stimulus recordings).

#### Dissection

Third instar larvae were washed and then dissected in ice-cold hemolymph saline (108 mM NaCl, 5 mM KCl, 2 mM CaCl_2_, 8.2 mM MgCl_2_, 4 mM NaHCO_3_, 1 mM NaH_2_PO_4_, 5 mM trehalose, 10 mM sucrose, and 5 mM Hepes at pH 7.5) ^10^. The anterior part of the larva was separated from the body just posterior to the mouth hooks. The head was then glued to a coverslip using tissue glue (3M, 084-1469SB). To make the antennal nerve accessible, the cuticle dorsal to the dorsal organ was removed. The cover slip with the dissected animal was placed into a flow chamber (Harvard Apparatus Inc., 640225) which was connected to two syringe pumps (World Precision Instruments, Aladdin2-220) for constant perfusion and to ensure smooth flow and evacuation of the odor. We estimate the turnover in the in chamber to be ∼15 s (chamber volume ∼500 μL, flow 33 μL/s).

#### Suction electrophysiology setup

Recording electrodes (World Precision Instruments, TW150F-3) were pulled (Sutter Instruments, P-87) and flame-polished (ALA Scientific Instruments, CPM-2). The electrode was connected to the head stage (Axon Instruments, CV-7B) of a microelectrode amplifier (Molecular Devices, Axon MultiClamp 700B) with a chlorinated silver wire. The head stage and electrode were placed on a motorized micromanipulator (Sutter Instruments, ROE-200 and MPC-200). The antennal nerve was sucked into the recording electrode by manually controlling external vacuum strength. The extracellular signal was amplified 100 times at the microelectrode amplifier, digitized (Molecular Devices, Digidata 1550B) and recorded at 10 kHz (Molecular Devices, Clampex Software). The signal was analog-filtered (10 kHz, Bessel) and 60-Hz main line noise was removed using an adaptive module in the digitizer (Molecular Devices, HumSilencer).

#### Odor delivery

At the electrophysiology rig, odors were delivered using multibarrel glass pipettes (World Precision Instruments Inc., 7B100F-4), which were pulled using a Micropipette puller (MicroData Instruments, PMP-107). One channel was filled with saline and the other channels were filled with the odor solution diluted in saline with the addition of fluoresceine (100 ng/μL; Sigma-Aldrich, F6377-100G) to visually assess the proper function of the odor pipette. An injection needle (Grainger, 5FVK8) was then inserted into the back of each barrel and sealed with hot glue. The channels were then connected to a multichannel pressure controller (Elveflow, OB1).

For the experiment, the tip of the odor stimulation pipette was placed and maintained directly next to the dorsal organ using a micromanipulator (Sutter Instruments, ROE-200 and MPC-200). To control the amount of odor presented during the experiment, pressure was varied in each channel at 50 Hz via previously published Python code (see ^33^ and *link to Zenodo upon publication*). As described elsewhere ^33,46^, odor flow resulted from the combination of the flow of one (i.e. Figure 1) or two (i.e. Figure 2) odor channel(s) and a channel carrying pure saline. The flow of the saline channel was adjusted to compensate for changes in the flow of the other channels such that the merged flow remained constant at a value of ∼1 nL/s (one odor channel) or ∼2 nL/s (two odor channels). The digitizer served as the timekeeper while the multichannel pressure controller reported each cycle (50 Hz).

#### Correlating odor concentrations in behavioral arrays and at the electrophysiology rig

In the behavioral experiments, animals are exposed to the odor in gaseous phase. The electrophysiology data were collected upon odor stimulation in liquid phase due to the immersion of the head preparation in saline. To correlate the concentration of the odor in gaseous phase with the concentration of odor in liquid phase, we made the following assumptions: (1) The diffusion of odor molecules into the structure of the dorsal organ is independent of the odorant receptor identity and, (2) the lowest concentration at which odor attraction was observed to airborne odor stimulation in the behavioral arena corresponds to the lowest concentration at which the spikes were observed in the electrophysiology recordings with odor stimulation in liquid phase.

Following this protocol, we identified the lowest amount of odor that animals were able to detect in the behavioral experiment and used our simulations of the odor diffusion (see section “**Physical model for odor diffusion**”) to calculate the average amount of odor in the air. At the electrophysiology rig, we investigated the lowest detectable concentration for each odorant. The liquid to air conversion rate was calculated as being the mean of the ratio of the detection threshold for airborne stimulation with simulated gaseous concentration (“Detection threshold gaseous”) divided by the detection threshold in liquid phase (“Detection threshold liquid”, see Table S3). Each odor was tested with both *Or42a* and *Or42b* OSNs. The final odor-specific liquid to air conversion factor was the mean of both ratios.

**Table S3:** In combination with the simulations of the time courses of the odor concentration experienced during behavior, this conversion factor was used to prepare the stimulus profiles of the trajectory replays shown in Figures 2 and 3.

| Odor | OSN | Detection threshold gaseous | Simulated gaseous concentration in air detection threshold $\theta_{air}$ | Detection threshold liquid $\theta_{liquid}$ | Liquid-to-air conversion factor $\theta_{air}/\theta_{liquid}$ | Odor specific liquid to air conversion factor $\langle \theta_{air}/\theta_{liquid} \rangle$ |
| --- | --- | --- | --- | --- | --- | --- |
| EtA | *Or42a* | 1:1,000 | 2.39e-7 M | 5e-6 M | 20.9 | 11.80 |
| EtA | *Or42b* | 1:1,000,000 | 3.6e-10 M | 1e-9 M | 2.7 |  |
| EtB | *Or42a* | 1:10,000 | 6.4e-9 M | 5e-8 M | 7.79 | 12.77 |
| EtB | *Or42b* | 1:100,000 | 5.6e-10 M | 1e-8 M | 17.7 |  |
| MA | *Or42a* | 1:100 | 4.3e-6 M | 1e-5 M | 23.07 | 12.12 |
| MA | *Or42b* | 1:100 | 4.3e-6 M | 5e-6 M | 1.15 |  |

#### Optogenetic stimulation

To activate the red-shifted light-gated ion channel ChrimsonR ^71^, a light-emitting diode (LED) at 625 nm (Mightex, PLS-0625-030-S) was mounted in the light path of the electrophysiology microscope (Olympus, BX51), which permitted consistent light stimulation between preparations through the 40x water immersion objective. Light intensity was adjusted with a LED controller (Mightex, SLA-1200-2), which was controlled by the digitizer.

#### Semi-automated manual spike sorting

Spike sorting and peristimulus time histogram (PSTH) analysis was performed as described in ref. ^33^ using custom Python software [*link to Zenodo upon publication*]. While *Orco* null mutants (O*rco^-/-^*) showed markedly decreased spontaneous activity in the antennal nerve, it was necessary to perform spike sorting on all recordings. We used a previously developed spike sorting algorithm ^33^ to aid manual annotation by a trained expert (D.T.). Briefly, the algorithm extracts vectors of different features of spikes (e.g., spike duration, spike amplitude). Uniform Manifold Approximation and Projection (UMAP) was then applied followed by hierarchical clustering using the HDBSCAN algorithm ^72^. A spike cluster of interest was then selected and manually checked for accuracy.

#### Analysis of electrophysiology data

PSTH was calculated as previously described ^33,46^. Briefly, each spike was convolved with a Gaussian filter, followed by grouping in 33 ms bins and averaged across experiments to yield the mean PSTH. Last, a Gaussian filter was applied to smoothen the signal. For the dose response curves (e.g. Figure 1D), for the phasic response to the odor stimulus was calculated as the mean firing rate over the first 2 s after stimulus onset. For the tonic response, the mean firing rate over the last 2 seconds of the 20 s stimulus was calculated.

#### Data exclusion procedure

All collected behavioral data are shown in the figures of the manuscript. The electrophysiological recordings were subject to several sources of variability that could not be controlled fully during data collection as described in ref. ^33^. Briefly, infrequent head movement due to twitching of the pharyngeal muscles could result in a temporary misalignment of the odor pipette relative to the animal with the resulting decrease in perceived odor concentration pushing the neuron out of depolarization block. To identify such trials, we screened recordings with the following fixed set of criteria: (i) sudden changes in local field potentials and (ii) delayed or missing responses during a control odor stimulus presented at the end of the recording. In addition, when the microfluidics controller delivering the odor was below 5% (10 mbar) of the maximum (200mbar) we noticed unreliable odor delivery. We therefore ignored these datapoints (Figures 2 and 3).

### Statistics

#### Statistical tests

Unless indicated otherwise, all datapoints were collected from different samples. No statistical methods were used to systematically predetermine sample sizes of each experimental condition. Custom-written Python scripts were used to create the plots and calculate the statistics can be downloaded through this link: [*link to Zenodo upon publication*].

To establish whether a parametric test could be used, we first tested for equality of variances with the Levene’s test (scipy.stats.levene with standard settings) and for normality of each sample using the Lilliefors test (statsmodels.stats.diagnostic.lilliefors with standard settings). To compare two groups, either the *t* test (scipy.stats.ttest_ind with standard settings) or the nonparametric Wilcoxon rank-sum test (scipy.stats.ranksums with standard settings) was used. To compare to baseline, either 0 (e.g. Figure 2E) or 0.5 (e.g. Figure 2J), either the one-sample *t* test (scipy.stats.ttest_1samp with standard settings) or the nonparametric Wilcoxon signed-rank test (scipy.stats.wilcoxon) was used.

All statistical tests were two-sided. To correct for multiple comparisons, Holm-Bonferroni multiple-test correction was used throughout the manuscript. To compare multiple groups, either ANOVA (scipy.stats.f_oneway with standard settings) followed by Tukey’s test (scipy.stats.tukey_hsd with standard settings) or the non-parametric Kruskal-Wallis test (scipy.stats.kruskal with standard settings) followed by the Conover-Iman test (scikit_posthocs.posthoc_conover with standard settings) was used. For all datasets, the sample size, mean, median, STD, SEM, 95% confidence intervals and effect size metrics (Hedges’ g, Cohen’s d, η and r, as described in ref. ^73^) can be found in supplementary file Data S1. For each test, the test statistic (such as the t-statistic for *t* tests), degrees of freedom, and exact p-values can be found in Data S1. Throughout the article, box plots contain the following elements: The orange center line indicates the median, the box includes the first (Q1) and third (Q3) quartiles, with the whiskers extending to Q1-1.5×IQR and Q3+1.5×IQR where IQR denotes the interquartile range. The notched part of the boxplot indicates the confidence interval around the median. All datapoints, including outliers, are plotted.

### Behavioral assay

The behavior apparatus consisted of round Petri dishes (Fisher Scientific, FB0875713) filled either with 25 mL 1% (w/v) agarose (Genesee Scientific, 20-102) or 1% (w/v) agarose + 36% (w/v) fructose (Millipore Sigma, F0127-1KG). Two types of odor cups were used: 3D printed single use odor cups (Figures 1-4 and corresponding supplementary figures) and reusable Teflon cups ^48^ (Figures 5-6 and corresponding supplementary figures). The 3D printed single use odor cups [*datadryad repository link to be added upon publication*] were produced with a Formlabs (Form 2) 3D printer, then washed and cured as instructed by the manufacturer.

Odors were diluted as indicated in paraffin oil (Sigma-Aldrich, 18512-1L) in glass vials with Teflon caps (Neta Scientific, 5182-0556). Odor dilutions were prepared daily. After vortexing for 10 seconds, 10 µl of diluted odor (or pure solvent in the ‘no odor’ condition) was placed into the odor cups immediately prior to each experiment.

Behavioral experiments in Figures 1-5 and corresponding supplementary figures were conducted using the PiVR tracker ^33,74^, which allowed video collection for *post-hoc* analysis of larval positions. Throughout all experiments, animals were tested only once (no repeated trials with the same larva). Petri dishes were never re-used.

#### Innate attraction assay

1% (w/v) agarose (Genesee Scientific, 20-102) was poured into round petri dishes (Fisher Scientific, FB0875713) and dried for approximately 30 min. Odor cups containing either the test odor at the indicated odor concentration or pure solvent were placed 5 mm away from one edge of the dish and an odor cup with solvent the same distance on the opposite edge of the dish. After 30 s, approximately 20 larvae of the indicated genotype were placed in the center of the dish, equidistant from the two cups. After 5 min, the number of larvae on the odor side, in the center (defined as a 7 mm-wide central zone), and on the solvent side were counted. Following an established protocol ^75^, the attraction index is calculated as (#odor-#no_odor)/#total. In this formula, # indicates the number of larvae on the corresponding side of the dish. The values of the *attraction index* range from -1 (total aversion to the test odor) to 1 (total preference for the test odor).

#### Equal naïve preference

For olfactory learning experiments, it is common to adjust the odor concentration such that naïve animals show equal attraction to both odors ^47,76^. We established the *equal naïve preference* by screening different odor combinations: one odor (e.g., EtA 1:50) was placed on one side of the dish and a second odor (e.g., EtB 1:200) on the opposite side of the dish. Approximately 20 larvae of the tested genotype were placed in the center of the dish. After 5 min, the number of larvae on both sides of the dish was counted. The Attraction Index was calculated as: (#odor1-#odor2)/#total. If a significant difference in attraction between the two odors was found, odor concentrations were adjusted to achieve an *attraction index* of 0 (see, for example, Figure 1-S1E).

#### Odor discrimination assay

To test whether larvae can discriminate between two odors, we adapted a learning assay described in detail in ref. ^75^. Groups of approximately 20 larvae were placed in a petri dish filled with 1% (w/v) agarose containing 36% (w/v) fructose and exposed to one odor (e.g., EtA 1:25) for 5 min. The animals were then moved to another dish without fructose and exposed to the second odor (e.g., EtB 1:400) for 5 min. This training cycle was repeated twice more for a total of three training cycles. For testing, the dish was filled with 1% (w/v) agarose without fructose, and the two odors (e.g., EtA 1:25 and EtB 1:400) were placed on opposite sides. The larvae were placed in the center of the dish, and after 3 min, the number of animals on each side was counted. Critically, each experiment included a corresponding reciprocal experiment run simultaneously: in the reciprocal experiment of the example above, a separate group of larvae was first exposed to EtB 1:400 with fructose, then to EtA 1:25 without fructose (repeated three times).

The analysis of the behavior proceeded in two steps: (a) the *preference index* was calculated for each experiment by subtracting the number of animals on the right side from those on the left side of the dish divided by the total number of animals: (#left-#right)/#total (e.g., Figure 1-S1H, left plot). (b) The *discrimination index* was then calculated from reciprocal pairs of experiments as (*preference index 1* – *preference index 2*)/2. This difference in preference between reciprocally trained groups reflects the ability to discriminate between the two tested odors. The reciprocal pairs used to calculate the *discrimination index* are indicated in the *preference index* plots by connecting lines (see, for example, Figure 1-S1H, left plot). A *discrimination index* significantly different from zero indicates that the two odors can be discriminated.

#### Odor identification assay

When the *discrimination index* is not significantly different from zero, several interpretations are possible: (1) the tested genotype cannot form an association between the odor and the fructose reward due to defects in memory formation; (2) the tested genotype cannot retrieve a successfully formed association due to defects in memory retrieval; (3) the tested genotype cannot discriminate between the odors due to limitations in how the olfactory sensory system encodes odors.

To rule out interpretation (1) and (2), we performed an *odor identification assay* for all genotypes. This assay tests the ability of each genotype to form and retrieve associations between tested odors and fructose. The procedure is identical to the *odor discrimination assay*, except that paraffin oil (the solvent) is used instead of a second odor. For example, in Figure 1-S1J, one group of larvae learned that EtA at 1:400 dilution is predictive of a fructose reward (left boxplot with the majority of larvae on the EtA side). The reciprocally trained group learned that the absence of odor predicts fructose (middle boxplot with the majority of larvae on the ‘no odor’ side). The *identification index* was calculated analogously to the *discrimination index*: first the *preference index* was calculated as (#left-#right)/#total, then the *identification index* was calculated as (*preference index 1* – *preference index 2*)/2. An *identification index* significantly different from zero indicates that the tested genotype can associate the tested odor with reward and retrieve this association, thereby ruling out interpretation (1) and (2).

### Computational model of OSN firing activity

#### Model of olfactory sensory neuron spiking activity

In our previous OSN model ^33^, the odorant transduction module describes the interactions of a single odor, ethyl butyrate, with the *Or42b*-expressing OSN. In this study, we adopt the same methodology to model the odorant transduction process. As before, we model the diffusion and absorption of odorant molecules through the sensilla of the OSN as follows:

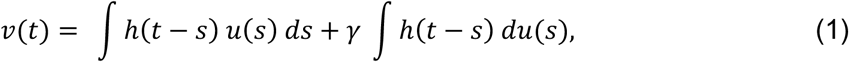

where 𝑢(𝑡) is the temporal waveform of the odor concentration experienced by the larva, ℎ(𝑡) is a low-pass linear filter, and 𝛾 is a weighting factor that determines the dependency of the filtered waveform 𝑣(𝑡) on the odor concentration 𝑢(𝑡) and the odor gradient 𝑑𝑢/𝑑𝑡. The output 𝑣(𝑡), referred to as the odorant concentration profile, then interacts with the private odorant receptors within the membrane of the OSN.

In the behavioral assay described here, there is an additional layer of complexity: the odor mixture interacts simultaneously with both the *Or42a* and *Or42b* OSNs. This process can be described using a competitive binding model, where different types of odorant molecules compete to bind to a limited number of receptors ^77,78^. This model assumes that only one odorant molecule can bind to a receptor binding site at a time, and that different odorant-receptor interactions have different responses in their binding and unbinding kinetics. Since we tested the responses of two OSNs (*Or42a*, *Or42b*) to three different odors (ethyl acetate, ethyl butyrate, and methyl acetate), there were a total of six combinations of odorant-receptor interactions to consider. By introducing competitive binding into the odor transduction model to account for odor mixtures, we have the following modified set of equations:

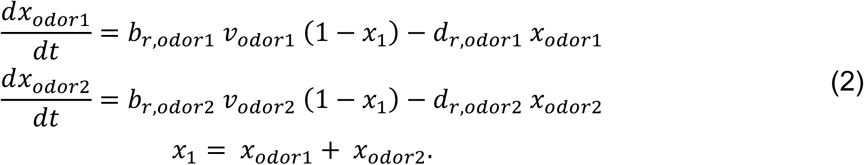

The variable 𝑥_1_ models the fraction of odorant receptors that are bound at any given time. Since we presented odor mixtures of two different odors, odorant receptors can be bound to the first odor, the second odor, or remain unbound. Therefore, by defining 𝑥_𝑜𝑑𝑜𝑟1_ and 𝑥_𝑜𝑑𝑜𝑟2_ as the fraction of odorant receptors bound to the two respective odors, we have: 𝑥_1_ = 𝑥_𝑜𝑑𝑜𝑟1_ + 𝑥_𝑜𝑑𝑜𝑟2_. The (filtered) odorant concentration profiles for each odor experienced by a larva are given by 𝑣_𝑜𝑑𝑜𝑟1_(𝑡) and 𝑣_𝑜𝑑𝑜𝑟_2(𝑡). Similarly, 𝑏_𝑟,𝑜𝑑𝑜𝑟1_ and 𝑏_𝑟,𝑜𝑑𝑜𝑟2_ define their respective association rate constants (see Table S4), while 𝑑_𝑟,𝑜𝑑𝑜𝑟1_ and 𝑑_𝑟,𝑜𝑑𝑜𝑟2_ define their respective dissociation rate constants. In the parameter fitting process, it was sufficient to vary only the association rates and treat the dissociation rates as constants. The association rates for each odor and OSN combination were optimized and are listed in Table S4 below:

**Table S4:** Optimized association and dissociation rate constants 𝑏_𝑟,𝑜𝑑𝑜𝑟_ and d_𝑟,𝑜𝑑𝑜𝑟_ of the odorant transduction model.

| OSN | Odor | $b_{r,odor}$ (ppm<sup>-1</sup> s<sup>-1</sup>) | $d_{r,odor}$ (s<sup>-1</sup>) |
| --- | --- | --- | --- |
| **Or42a** | Ethyl acetate | $2.18 \times 10^{-1}$ | 10 |
| **Or42a** | Ethyl butyrate | $2.17 \times 10^{-2}$ | 10 |
| **Or42a** | Methyl acetate | $3.10 \times 10^{-4}$ | 10 |
| **Or42b** | Ethyl acetate | $6.04 \times 10^{-5}$ | 10 |
| **Or42b** | Ethyl butyrate | $5.30 \times 10^{-3}$ | 10 |
| **Or42b** | Methyl acetate | $1.38 \times 10^{-5}$ | 10 |

For the modelling of the co-receptor channel 𝑥_2_, and the calcium channel 𝑥_3_, we adopt the following equations:

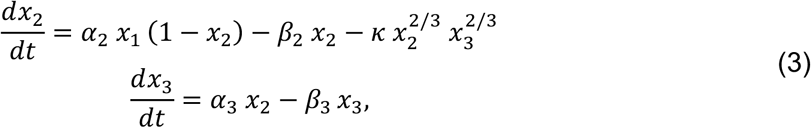

where 𝑥_1_ is the total fraction of odorant receptors bound by both odors. The parameters of eq. (3) are defined in Table S5.

**Table S5:** Optimized parameters of the odor transduction model. The input of the odor transduction model is the gaseous odor concentration 𝑢 (in ppm) experienced by the animals, and the output is the transduction current 𝐼 (in μA cm^-2^). To convert the odor concentration 𝑢 delivered through the odor pipette in liquid phase (measured in µM) into the equivalent odor concentration that would be experienced by the animals in gaseous phase (in ppm), the odor-specific liquid to air conversion factor in Table S3 was applied. **[*]** indicates parameter values different from ref. ^79^.

| Parameter | Physical description | Parameter value(s) | Units |
| --- | --- | --- | --- |
| **h(t) [\*]** | Low-pass linear filter modeling the peri-receptor process, defined as the solution to the differential equation:<br>$\frac{d^2}{dt^2} h(t) + 2 \alpha_1 \beta_1 \frac{d}{dt} h(t) + \alpha_1^2 h(t) = \alpha_1^2 \delta(t)$<br>($\delta(t)$ is the Dirac delta function) | $\alpha_1 = 29.3$<br>$\beta_1 = 14.7$ | s<sup>-1</sup><br>unit-less |
| **$\alpha_3$ [\*]** | Rate of increase of the calcium gating variable | 8.508 | s<sup>-1</sup> |
| **$\beta_3$ [\*]** | Rate of decrease of the calcium gating variable | 0.296 | s<sup>-1</sup> |
| **$\gamma$ [\*]** | Scaling factor of the filtered odorant concentration gradient | 1.5 | s |
| $\alpha_2$ | Rate of activation of the co-receptor channel gating variable | 146.1 | s<sup>-1</sup> |
| $\beta_2$ | Rate of deactivation of the co-receptor channel gating variable | 117.2 | s<sup>-1</sup> |
| $\kappa$ | Feedback strength from the calcium channel to the c-receptor channel | 8841 | s<sup>-1</sup> |
| $c$ | Half-activation coefficient of the co-receptor channel | $6.546 \times 10^{-2}$ | unit-less |
| $p$ | Hill coefficient of the co-receptor channel | 1 | unit-less |
| $I_{max}$ | Maximum transduction current generated by the co-receptor channel | 62.13 | $\mu\text{A cm}^{-2}$ |

All other equations in the odorant transduction model are identical to those in our previous work. Note that besides the odorant-receptor specific association/dissociation rates added to the odor transduction model, all other parameters remain the same as found in our previous OSN model ^33^. The current resulting from this odorant transduction process is then described as a Hill function of the co-receptor gating variable 𝑥_2_:

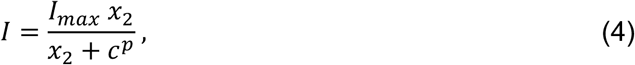

where 𝐼*_max_* is a time-dependent constant that defines the maximum transduction current that can result through the co-receptor channel. The parameters 𝑐 and 𝑝 define the half-activation coefficient and Hill coefficient of the co-receptor channel, respectively.

The spike generation component of the OSN model is based on the 3-element model of Qian et al. ^80^, which contains three state variables: 𝑉, ℎ, and ℎ*_s_*. The feature of the Qian model that differs from other point neuron models is the distinction between fast inactivation (ℎ) and slow inactivation (ℎ*_s_*) in the sodium current, which captures the dynamics of depolarization block.

The Qian 3D model consists of the following system of ordinary differential equations:

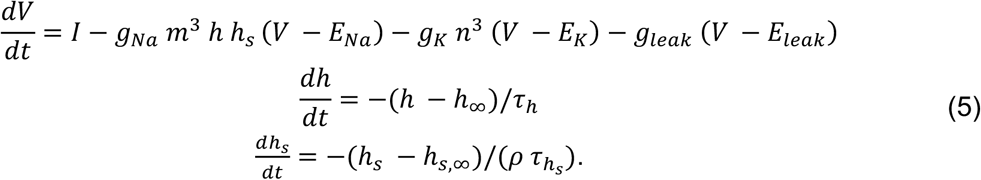

In the above equations, 𝐸*_Na_*, 𝐸*_K_* and 𝐸*_leak_* are the respective Nernst potentials of each channel. Parameters 𝑔*_Na_*, 𝑔*_K_* and 𝑔*_leak_* model the maximal conductance per unit area of each channel. The symbols 𝑛, 𝑚, ℎ, and ℎ*_s_* are dimensionless state variables mediating the gating of the three currents, modelled as time-dependent functions of the membrane potential 𝑉. Parameters ℎ_∞_ and ℎ*_s_*_,∞_ are voltage-dependent steady-state values of the fast and slow inactivation variables associated with the sodium channels. Parameters 𝜏*_h_* and 𝜏*_h_s__* are voltage-dependent time constants that modulate the rate of inactivation. For the spike generator model, we assumed that the firing dynamics were similar across OSNs besides the different responses in odorant transduction. Therefore, we adopted the same spike generator model for both the *Or42a* and *Or42b* OSNs, with identical parameters except for minor differences for the sodium channel. Table S6 below lists the differences between *Or42a* and *Or42b* OSN models:

**Table S6:** Parameter differences between the *Or42a* and *Or42b* OSN models.

| Parameter | Description | *Or42a* Value | *Or42b* Value | Units |
| --- | --- | --- | --- | --- |
| $g_{Na}$ | Conductance of the sodium channel | 14.4 | 18 | mS cm<sup>-2</sup> |
| $\rho$ | Scaling parameter for slow sodium channel time constant | 20 | 31 | unitless |

Parameters of the *Or42a* and *Or42b* OSN models were optimized using electrophysiology recordings for different stimulus patterns for each odor. This included step stimuli at different concentrations, along with linear, exponential, and sigmoidal ramp (Figures 3-S2 to 3-S5). Parameter optimization involved fitting the association/dissociation rates for each odorant-receptor pair using the global optimization toolbox in MATLAB with its in-built genetic algorithm. The objective was to minimize the normalized root mean squared error between the experimentally recorded firing rate and the model-predicted firing rate of the OSN. In the parameter optimization procedure, we matched the number of simulated trials with the existing number of experimental trials.

### FT-IR measurements for odor concentration profiles

To characterize the spatial distribution of EtA, EtB, and MA gradients, we used FT-IR measurements with a Tensor 27 (Bruker) following the protocol established in ref. ^43^. As illustrated in Figure 2-S1, two distinct configurations were used: one with the odor source positioned adjacent to the infrared light beam path (close) and another with the odor source located opposite to the infrared light beams (remote). To accurately model the odor diffusion within the behavioral arena, we fit the model parameters to match the FT-IR measurements obtained at source dilutions of 1/10 and 1/20 for EtB and MA, and at source dilutions of 1/10 and 1/50 for EtA. The raw absorption data and conversion into concentrations are provided in Data S2.

### Physical model for odor diffusion

The sensory experience of larvae in the behavioral assay was inferred using the same methodology outlined in ref. ^33^, using a 3D odor diffusion model. In contrast with the behavioral assay developed to study depolarization block ^33^, we used a circular arena instead of a rectangular arena, modelled with the open-source CAD software OpenSCAD ^81^. The arena had a diameter of 85 mm and a height of 7 mm, as illustrated in Figure 2-S1A. Experimentally, the main difference to the previous assay ^33^ was the use of cylindrical odor cups to contain the odor droplet rather than reinforcement rings. Due to the small aperture of the odor cups, the odor flux at the droplet-air boundary was smaller than that observed in our previous work, resulting in a more gradual change in the odor profile over time.

To simulate this difference, we modeled the geometry of the cylindrical odor cup with a radius of 3 mm and a height of 4 mm. The odor droplet has a volume of 𝑉_𝑑𝑟𝑜𝑝_ = 10 μL and is placed inside the odor cup in a cylindrical enclosure with a radius of 2 mm and a height of 0.8 mm. The walls of the odor cup are assumed to be no-flux boundaries, and the odor droplet diffuses out the top of the odor cup through a single aperture with a radius of 0.5 mm.

In each experiment, the odor cup is positioned 37.5 mm away from the center of the arena. We assume that the initial odorant concentration in the air is zero, while the initial odorant concentration in the droplet is equivalent to the applied source concentration. In simulations of different odor dilutions, we adjust the applied source concentration while holding all other aspects of the model constant. The parameters of the model were fit to match Fourier transform-infrared spectroscopy (FT-IR) measurements ^43^ captured along two different cross sections of the arena at different time intervals. In the “close” configuration, the FT-IR beam is positioned along an axis with an angular separation of 18° from the odor cup (Figure 2-S1A). In the “remote” configuration, the FT-IR beam is positioned further away with an angular separation of 90° from the odor cup.

For each odor used in the odor discrimination learning assay, FT-IR measurements were collected at the close and remote configurations using two different source dilutions. This resulted in different temporal odor profiles as shown in Figure 2-S1B-D for ethyl acetate, ethyl butyrate, and methyl acetate. To fit the diffusion parameters for each odor, we performed parameter optimization using the global optimization toolbox in MATLAB. The objective was to minimize the normalized root mean squared error (RMSE) between the experimental and simulated odor concentrations observed along the close and remote configurations. After parameter optimization, we found that our odor simulations (solid lines) were in excellent agreement with the experimental odor profiles (dashed lines) shown in Figure 2-S1B-D. In each simulation, we assumed a 30-s delay during which the odorant diffuses from the odor cup into the air before larvae are introduced into the arena. We also assumed, in the simulation of multiple odor sources, that there are no interactions between odorant molecules affecting the individual odor profiles (Figure 2-S1F).

### Trajectory reconstruction of individual animals from video recordings

For each experiment, we identified the positions of individual larvae during both the training and test phases. This was achieved using an adapted version of the motion-based multiple object tracking algorithm in MATLAB to automate the extraction of larval trajectories from video datasets (Figure 3-S1A). The *discrimination index* calculated using the automated tracking algorithm was found to be in excellent agreement with manually counted values (data not shown). In the original off-the-shelf implementation, the tracking algorithm performed well when larvae were well separated and away from the odor sources, but performed poorly when larvae aggregated near the odor sources. To avoid losing track of larvae at the odor sources, a counter was added to track larvae entering and leaving the vicinity of the odor sources.

### Plotting of calcium response of the complete olfactory system to EtA and EtB

Data S1 from Si et al. ^14^ *(Raw Activity Data of 21 ORNs Responding to 34 Odorants at Multiple Concentration Levels Collected from 238 Recordings)* was used to generate Figure 5-S1 which represents the mean dF/F across animals per concentration.

## Data and code availability

– All data underlying all figures will be deposited and made publicly available on Dryad (https://datadryad.com) upon publication.
– All original code will be deposited and made publicly available on Zenodo upon publication.
– Any additional information required to reanalyze the data reported in this paper is available from the lead contact upon request.

## Author contributions

DT performed all electrophysiology experiments and behavioral experiments in Figure 1. DT supervised behavioral experiments in Figure 4. NK performed behavioral experiments in Figure 4. MLM and MP performed behavioral experiments in Figures 5-6. PW performed odor diffusion and OSN modeling (Figure 3) and computational analysis of larval trajectories. JM supervised model development. ML designed the study, performed FT-IR measurements, and supervised the work and data interpretation. ML and DT wrote the manuscript with input from all authors.

---

**Figure 1-S1:**
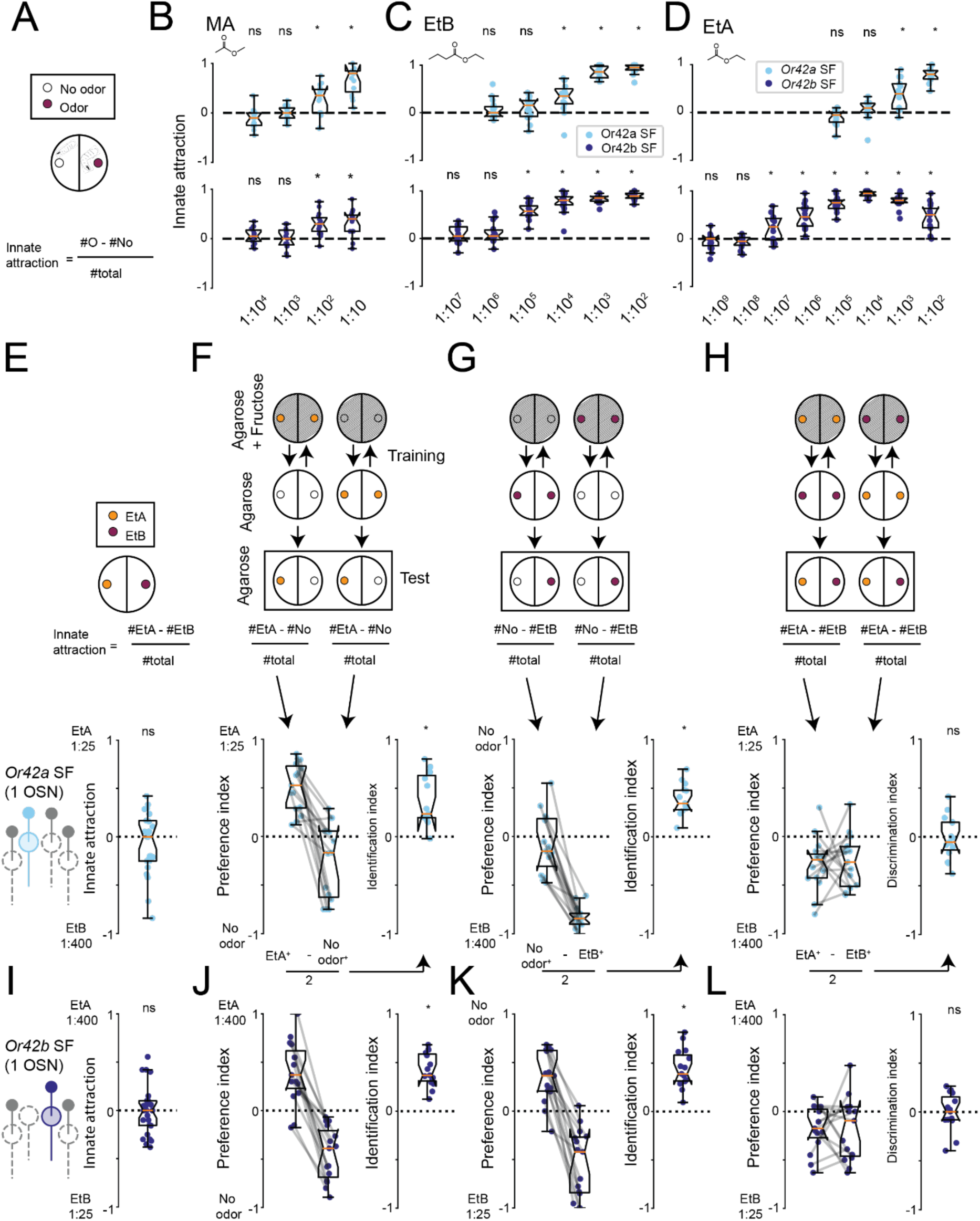
Single functional OSNs are insufficient for odor discrimination. **(A)** Schematic of the innate attraction assay. Larvae were placed in a dish with odor on one side and no odor on the other; the innate attraction index was calculated from the number of larvae on each side. **(B)** *Or42a* single functional (SF) and *Or42b* SF are attracted to MA at similar odor concentration ranges (n=15, one-sample *t* test followed by Holm-Bonferroni correction). **(C)** *Or42a* SF and *Or42b* SF larvae show largely overlapping sensitivity to EtB (*Or42a* SF: n=15, Wilcoxon signed-rank test; *Or42b* SF: n=15, one-sample *t* test; Holm-Bonferroni correction). **(D)** *Or42b* SF larvae are several orders of magnitude more sensitive to EtA than *Or42a* SF larvae (n=15, Wilcoxon signed-rank test followed by Holm-Bonferroni correction). **(E)** Naïve *Or42a* SF larvae show equal attraction to EtA 1:25 and to EtB 1:400, indicating similar detectability and valence (n=25, one-sample *t* test). **(F)** *Or42a* SF larvae associate both the presence and absence of EtA with a reward in the identification assay (see Methods). Gray lines connect reciprocal training conditions. *Or42a* SF larvae successfully learn EtA associations (n=15, Wilcoxon signed-rank test). **(G)** *Or42a* SF larvae associate EtB with reward (n=15, one-sample *t* test). **(H)** *Or42a* SF larvae fail to discriminate between EtA and EtB in the discrimination assay (n=15, one-sample *t* test). **(I)** *Or42b* SF larvae show equal attraction to EtA 1:400 and EtB 1:25 (n=26, one-sample *t* test). **(J,K)** *Or42b* SF larvae successfully associate presence and absence of EtA (n=15) and EtB (n=15) with reward (one-sample *t* test). **(L)** *Or42b* SF larvae fail to discriminate between EtA and EtB (n=15, one-sample *t* test). Throughout the figure, *p<0.05.

**Figure 1-S2:**
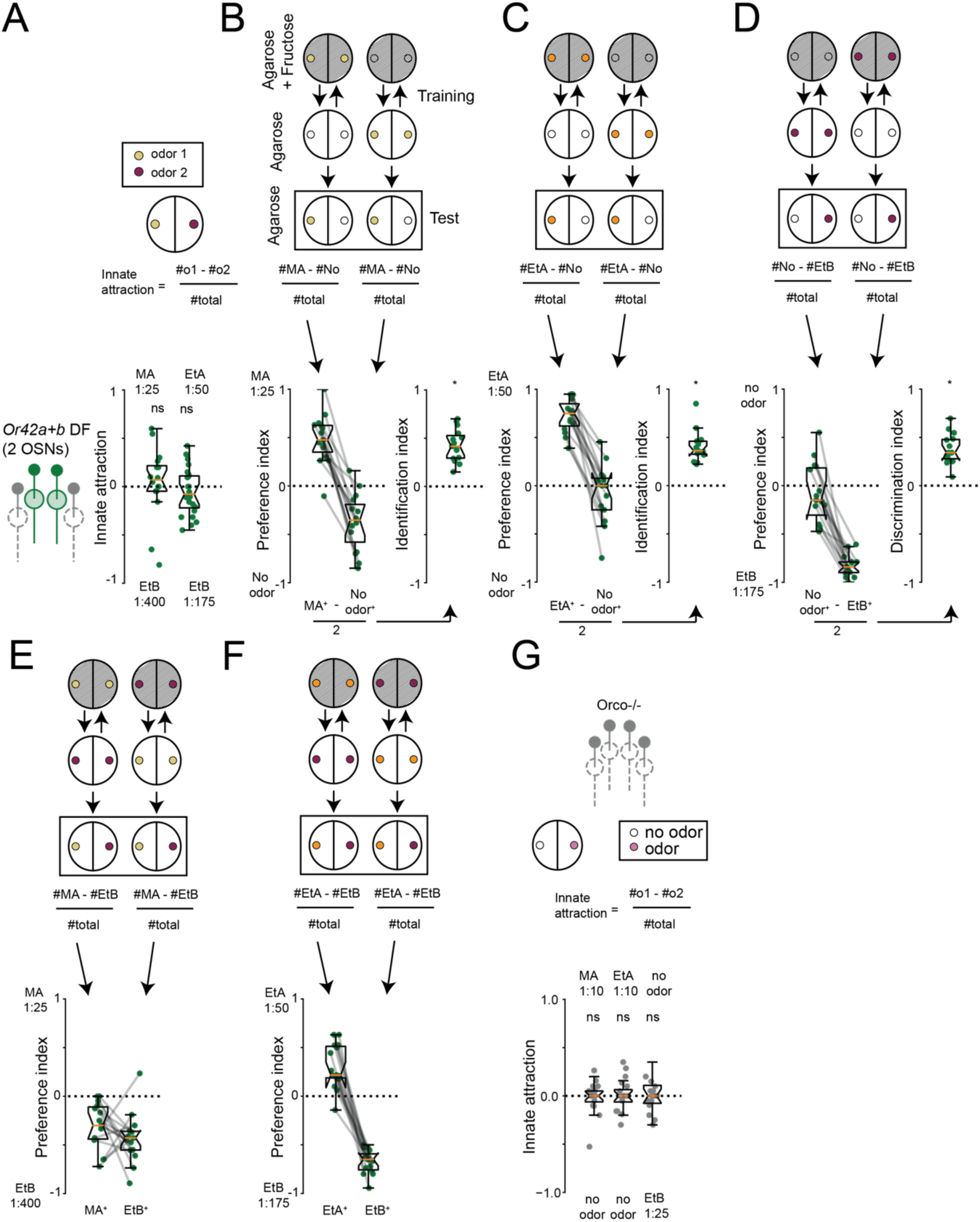
*Or42a+b* double functional (DF) larvae associate all tested odors with the reward. **(A)** *Or42a+b* DF larvae show equal attraction to MA 1:25 vs. EtB 1:400 (n=16) and to EtA 1:50 vs. EtB 1:175 (n=24, one-sample *t* test followed by Holm-Bonferroni correction). **(B-D)** *Or42a+b* DF larvae associate presence and absence of (B) MA, (C) EtB and (D) EtA with a reward (n=15 each, one sample *t*-test). **(E, F)** Reciprocal preference indices of *Or42a+b* DF larvae underlying discrimination results in Figure 1L (n=15, one-sample *t* test). **(G)** *Orco* null larvae cannot detect the highest odor concentrations tested (n=15, one-sample *t* test followed by Holm-Bonferroni correction). Throughout the figure, *p<0.05.

**Figure 2-S1:**
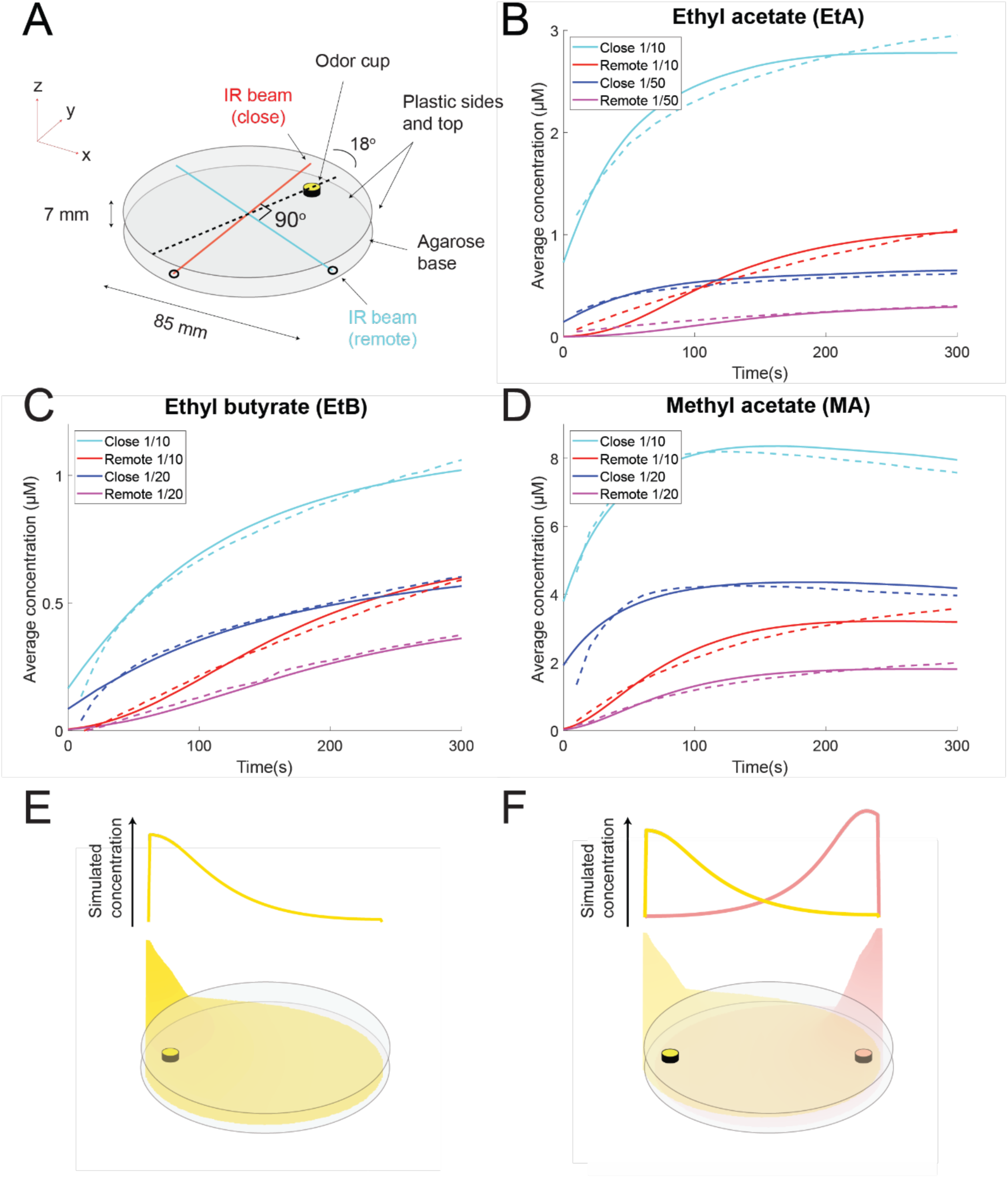
Odor diffusion model for discrimination learning assay. **(A)** Arena geometry and simulation features. Odor profiles were measured by FT-IR absorption along two cross-sections (close and remote) ^43^. **(B-D)** Comparison of FT-IR derived (dashed lines) and simulated (solid lines) odor concentration profiles for (B) methyl acetate, (C) ethyl butyrate, and (D) ethyl acetate. **(E)** Simulated odor profile for a single ethyl butyrate source. **(F)** Simulated profile for an odor mixture of ethyl butyrate (left) and ethyl acetate (right) sources. (n=3-4 independent measurements per odor, see Table S2).

**Figure 3-S1:**
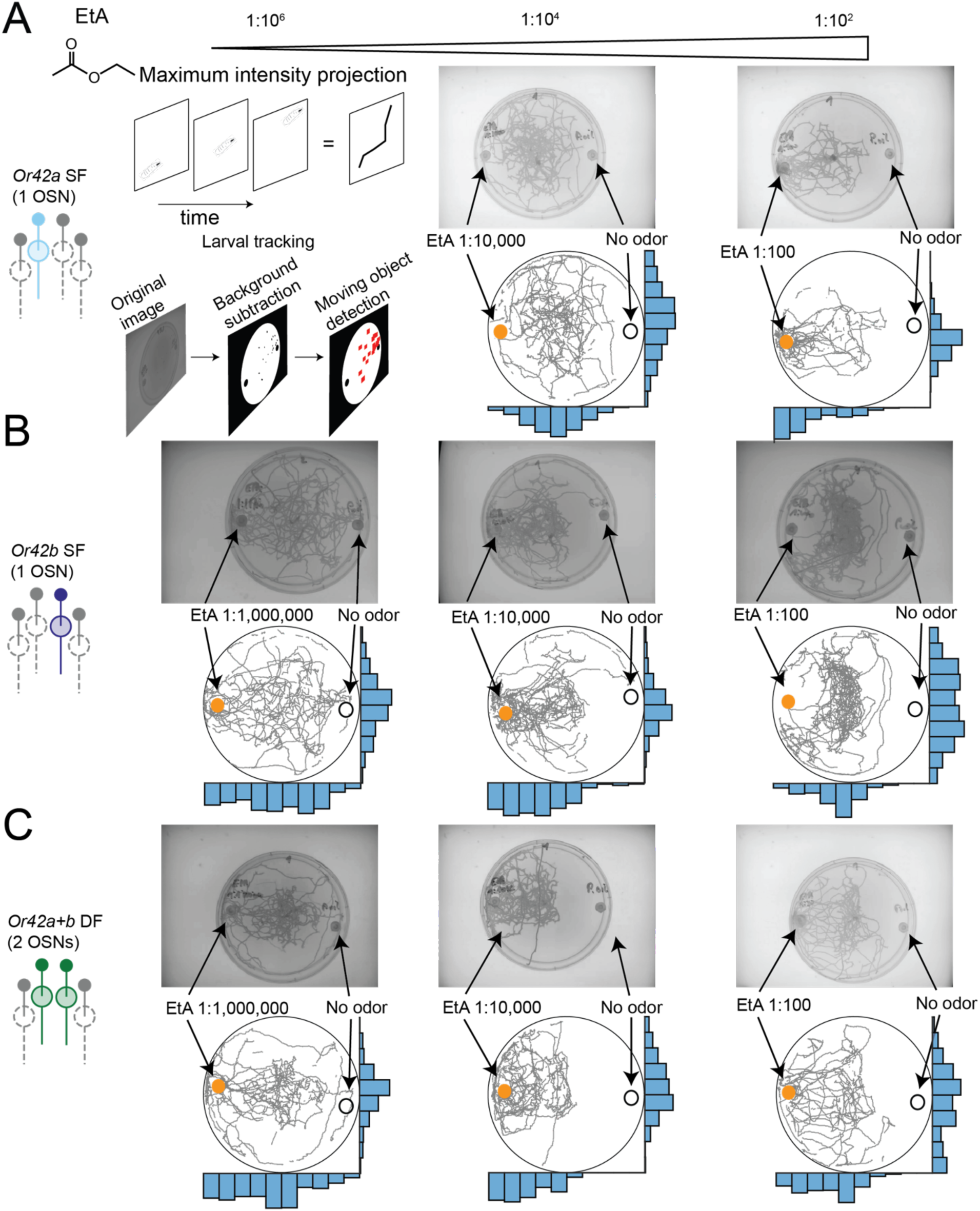
*Or42a+b* DF larvae are attracted over a wider concentration range than single functional larvae. Note: Trajectories and spatial distributions shown are uncorrected for dish tilt; tilt-corrected data were used for quantitative analyses in Figure 3. **(A)** Top row: Maximum intensity projection of videos. Bottom row: extracted larval trajectory segments using automated object tracking (see Methods). *Or42a* SF larvae are strongly attracted to high EtA concentrations. Histograms are calculated on trajectory segments that were automatically extracted from videos (top row: maximum intensity projection). **(B)** *Or42b* SF larvae are strongly attracted to EtA at 1:10,000. At higher concentrations (e.g., 1:100), most larvae avoid the odor source due to depolarization block ^33^. **(C)** *Or42a+b* DF larvae show attraction over a wider concentration range than single functional larvae.

**Figure 3-S2:**
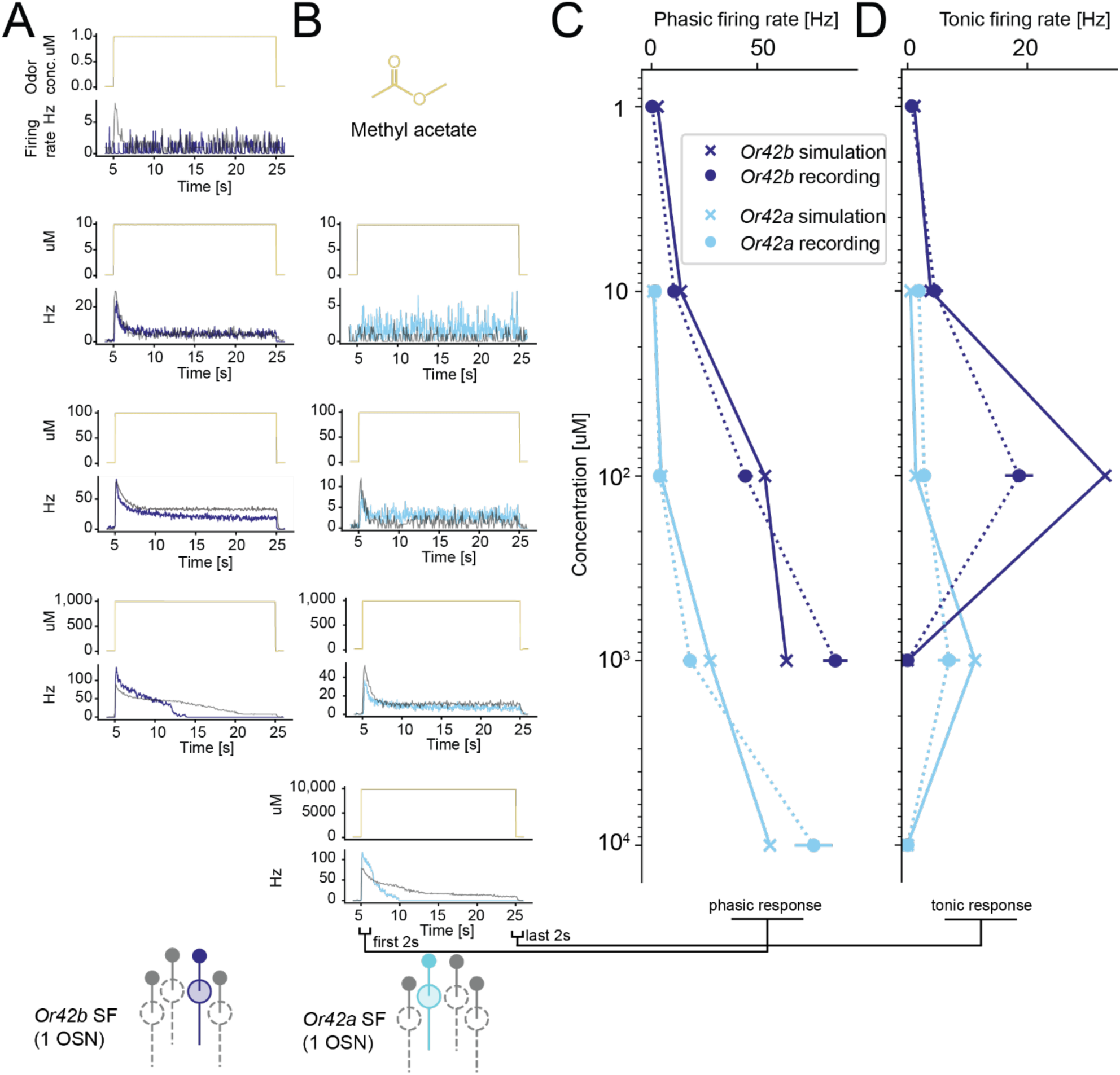
OSN model fitting with methyl acetate. **(A)** *Or42b* OSN (1 μM n=9, 10 μM n=18, 100 μM n=10, 1 mM n=9) and **(B)** *Or42a* OSN responses to 20-second step stimuli of methyl acetate at the indicated concentration (10 μM n=10, 100 μM n=19, 1 mM n=10, 10 mM n=10). Gray lines show model fits recapitulating depolarization block at higher concentrations. **(C)** Dose-response curve for *Or42a* and *Or42b* OSN phasic responses and **(D)** steady state (circles) and model fits (crosses).

**Figure 3-S3:**
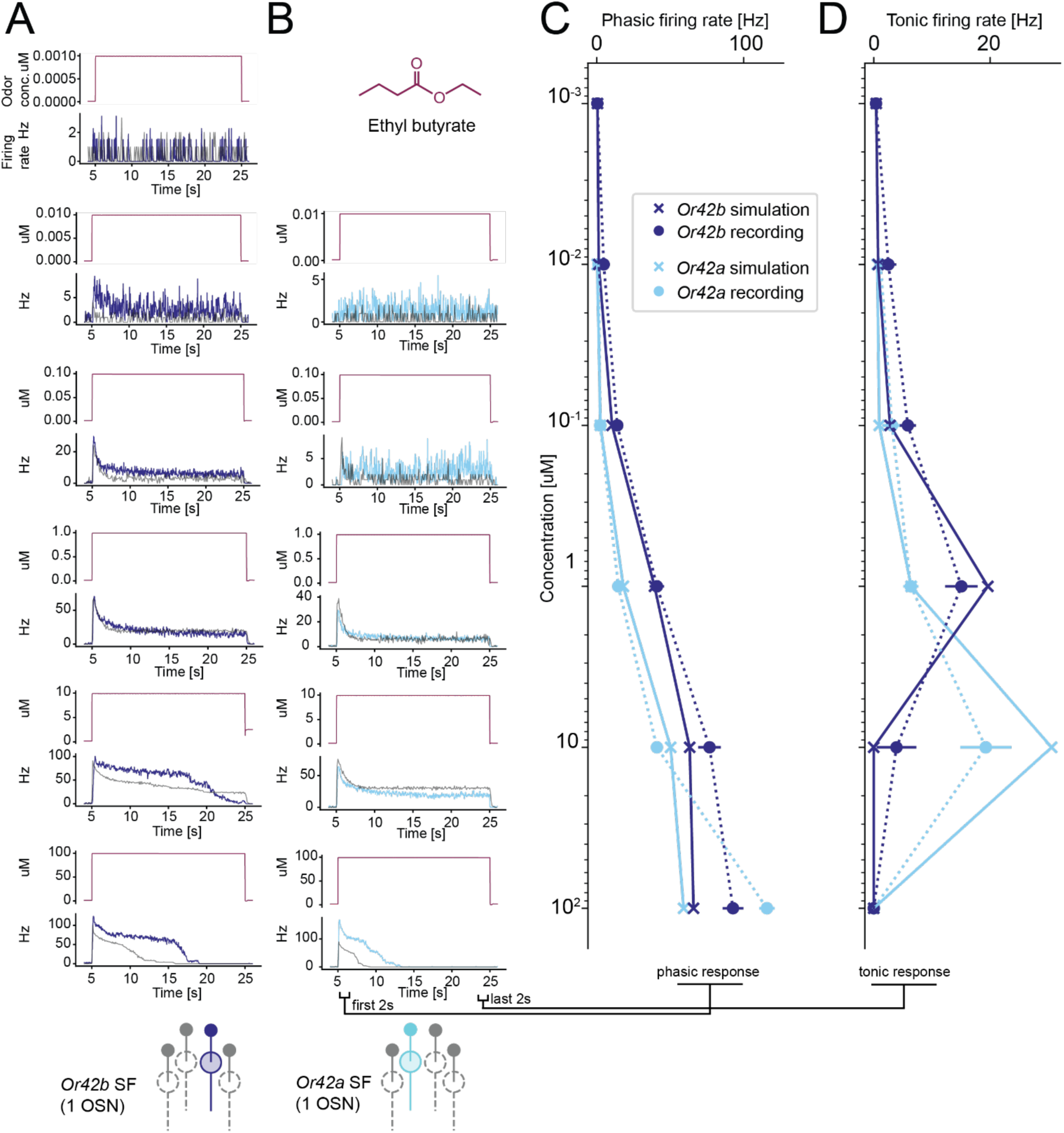
OSN model fitting with ethyl butyrate. **(A)** *Or42b* OSN (1 nM n=10, 10 nM n=10, 100 nM n=18, 1 μM n=10, 10 μM n=9, 100 μM n=10) and **(B)** *Or42a* OSN responses to 20-second step stimuli of ethyl butyrate (10 nM n=12, 100 nM n=10, 1 μM n=33, 10 μM n=14, 100 μM n=10). Gray lines show model fits recapitulating depolarization block at higher concentrations. **(C)** Dose-response curves for *Or42a* and *Or42b* OSN phasic responses and **(D)** steady state responses (circles) with model fits (crosses). Data in (A) from ref. ^33^.

**Figure 3-S4:**
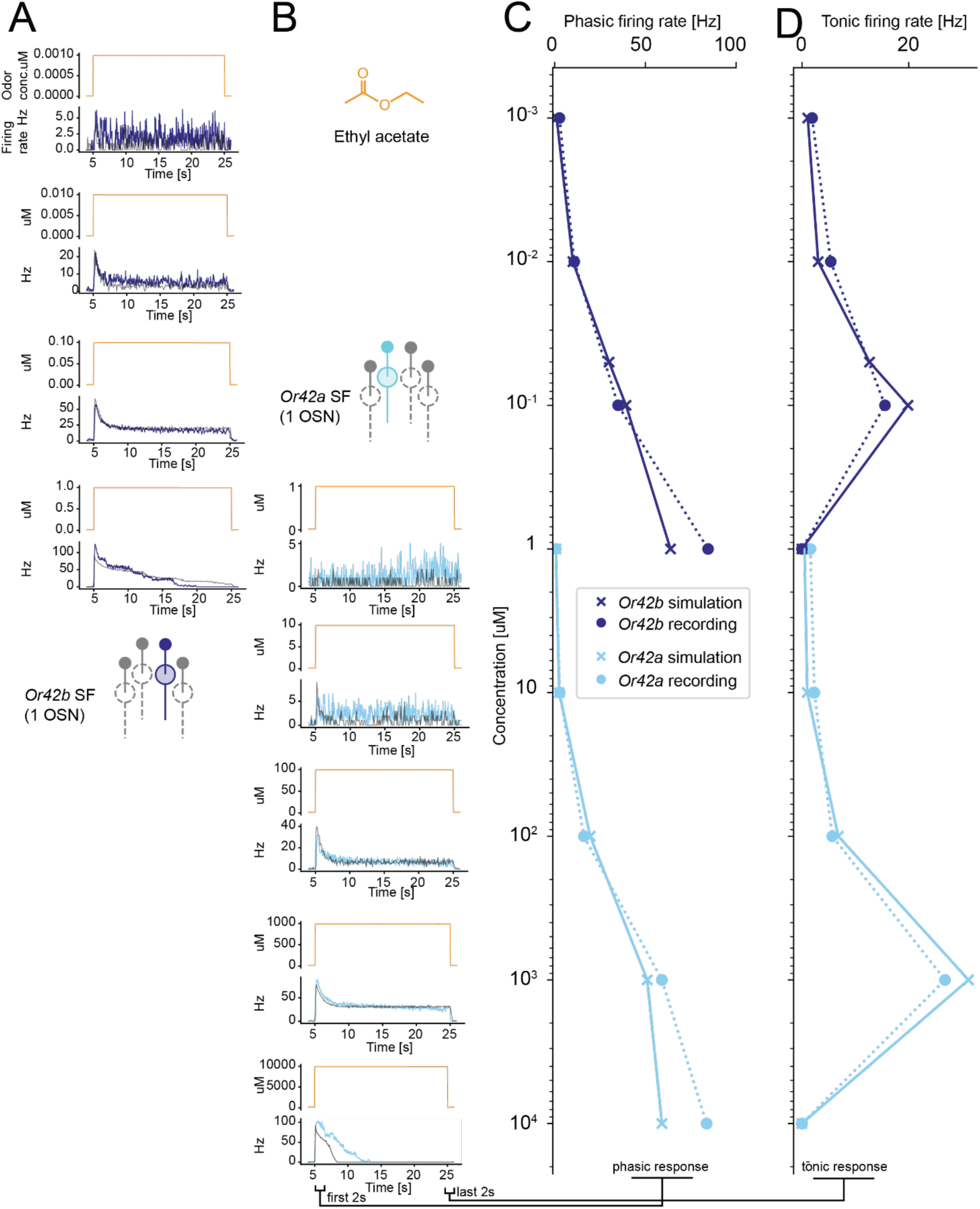
OSN model fitting with ethyl acetate. **(A)** *Or42b* OSN (1 nM n=10, 10 nM n=16, 100 nM n=22, 1 *μM* n=10) and **(B)** *Or42a* OSN responses to 20-second step stimuli of ethyl acetate (1 μM n=12, 10 μM n=14, 100 μM n=12, 1 mM n=9, 10 mM n=10). Gray lines show model fits recapitulating depolarization block at higher concentrations. **(C)** Dose-response curves for *Or42a* and *Or42b* OSN phasic responses and **(D)** steady-state responses (circles) with model fits (crosses).

**Figure 3-S5:**
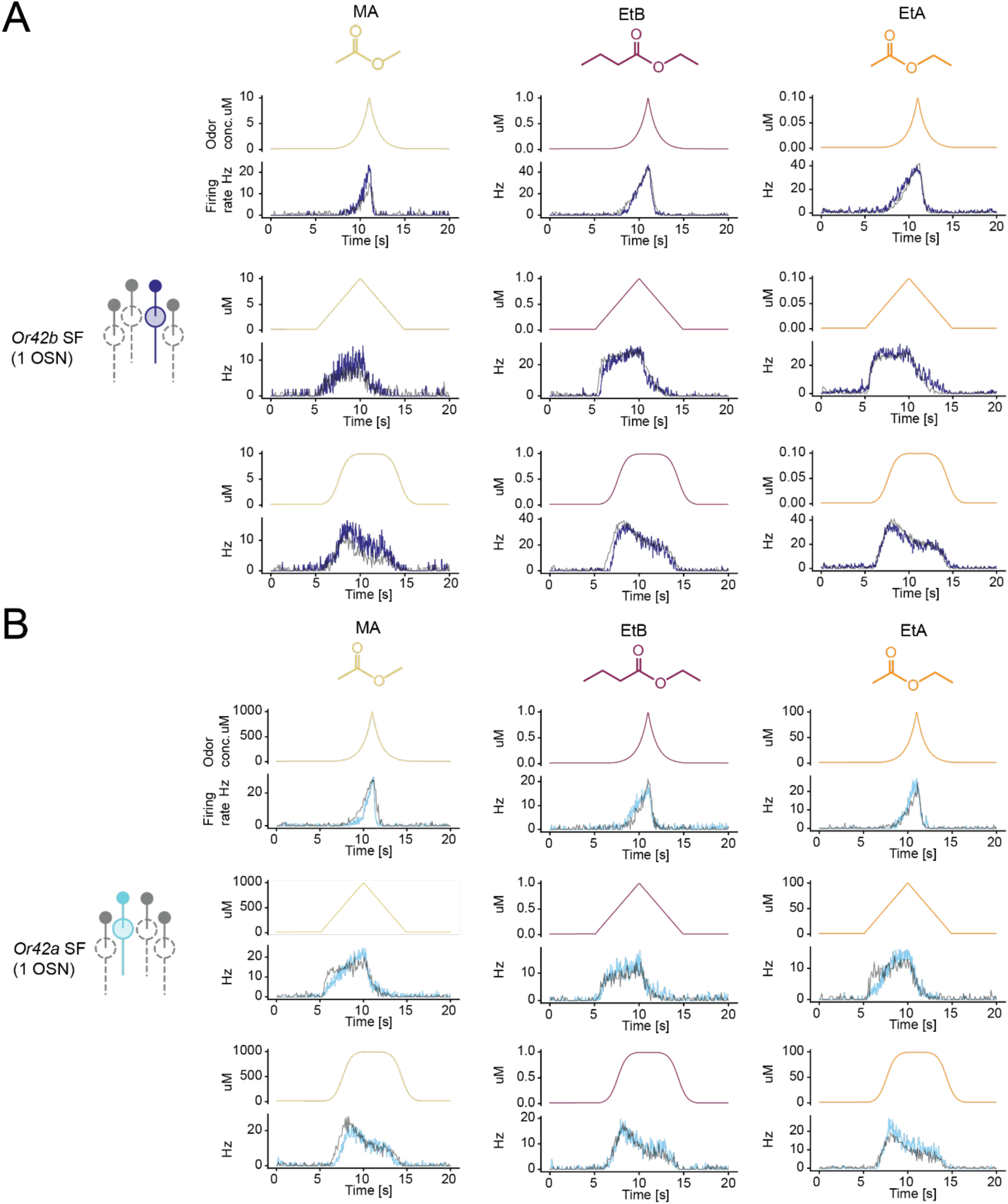
OSN model fitting with dynamic ramps. **(A)** *Or42b* OSN (exponential: MA n=8, EtB n=10, EtA n=12; linear: MA n=8, EtB n=10, EtA n=12; sigmoidal: MA n=8, EtB n=8, EtA n=12) and **(B)** *Or42a* OSN responses to dynamic ramps of methyl acetate (left), ethyl butyrate (center) and ethyl acetate (right) (exponential: MA n=10, EtA n=10, EtB n=12; linear: MA n=10, EtB n=12, EtA n=13; sigmoidal: MA n=10, EtB n=12, EtA n=10). Gray lines show model fits.

**Figure 4-S1:**
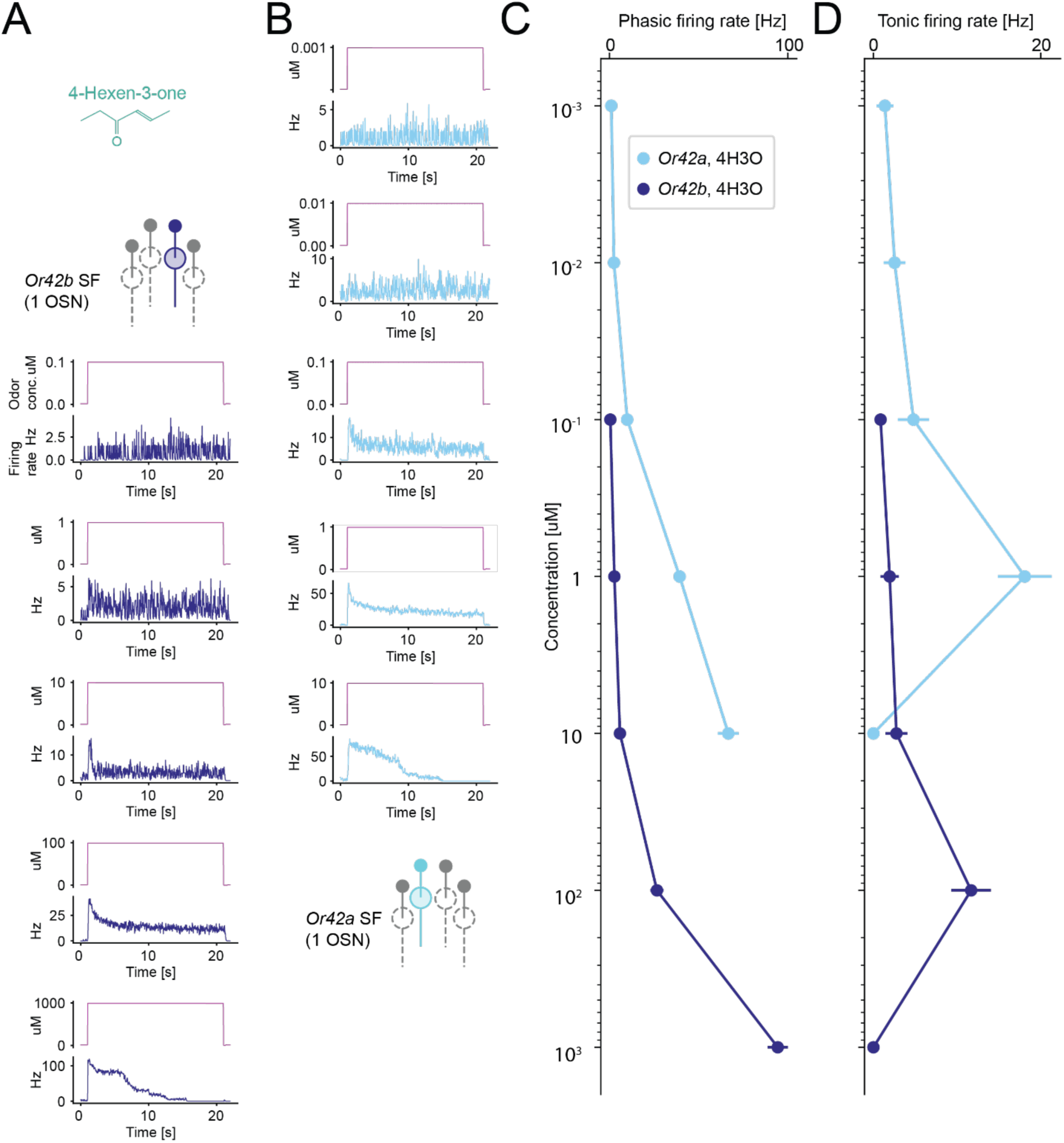
Electrophysiology recordings with 4-hexen-3-one. **(A)** *Or42b* OSN (100 nM n=10, 1 μM n=10, 10 μM n=10, 100 μM n=10, 1 mM n=8) and **(B)** *Or42a* OSN responses to 20-second step stimuli of 4-hexen-3-one (1 nM n=8, 10 nM n=10, 100 nM n=9, 1 μM n=10, 10 μM n=7). **(C)** Dose-response curves for *Or42a* and *Or42b* OSN phasic responses and **(D)** steady-state responses.

**Figure 4-S2:**
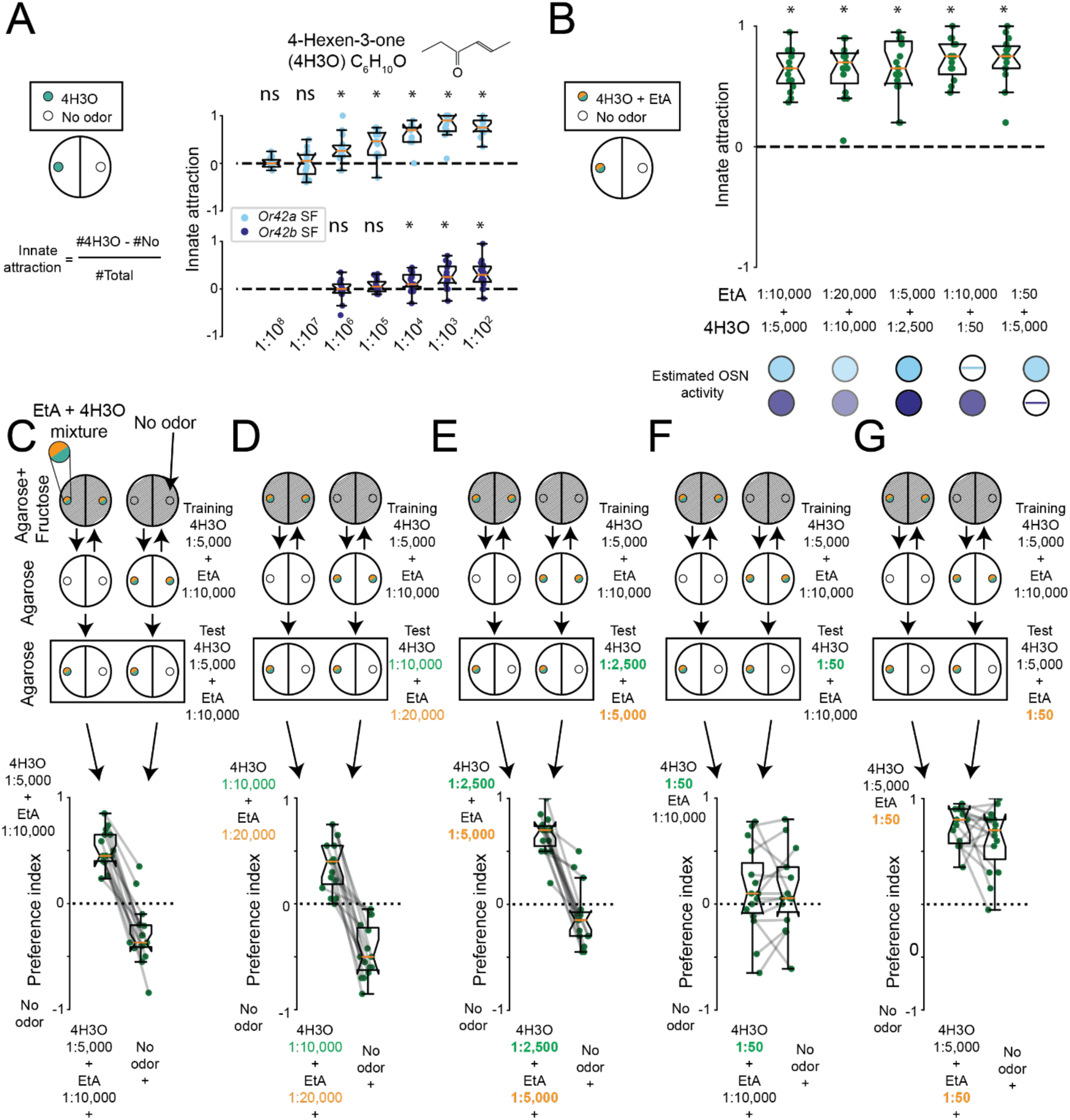
*Or42a+b* DF larvae show innate attraction to all tested odor mixtures. **(A)** Dose response curves for *Or42a* SF (top) and *Or42b* SF (bottom) to 4-hexen-3-one (4H3O) (n=15, one-sample *t* test followed by Holm Bonferroni correction). **(B)** All odor mixtures in Figure 4D-M elicit innate attraction (n=15, one-sample *t* test with Holm-Bonferroni correction). **(C-G)** Reciprocal preference indices underlying discrimination results in Figure 4E, G, I, J, and M, respectively. Sample sizes given in Figure 4.

**Figure 5-S1:**
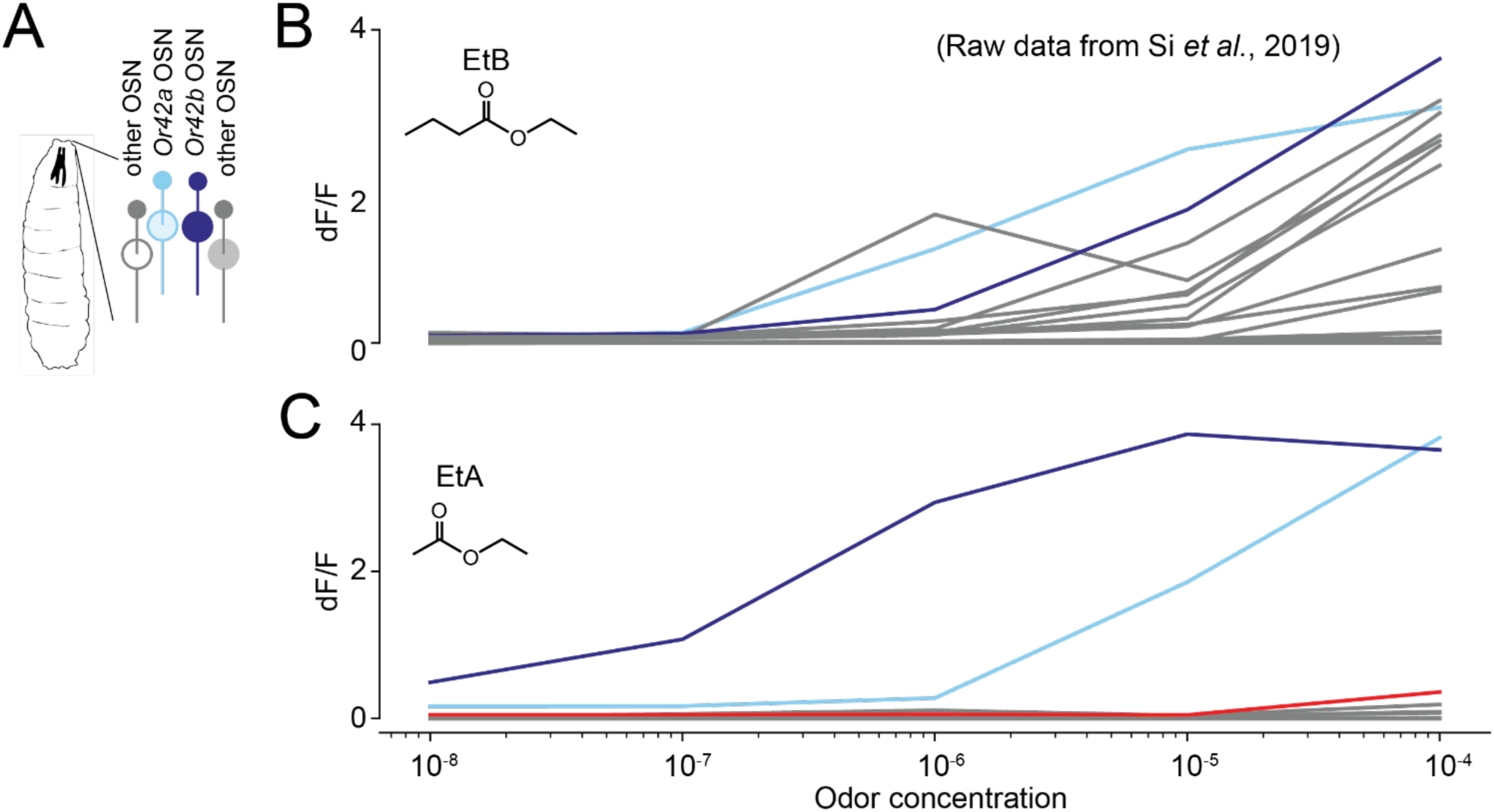
Calcium imaging of whole olfactory system responses to EtA and EtB. **(A)** Schematic of larval olfactory sensory system (data from ref. ^14^). **(B)** *Or42a* (cyan), *Or42b* (navy) and other OSNs have similar phasic mean dose-response curves to ethyl butyrate during 5-second stimulation. **(C)** During 5-second step ethyl acetate stimulation, *Or42b* OSN is most sensitive, followed by *Or42a* OSN. Other OSNs including *Or33a* and *Or47a* (red) respond only at the highest EtA concentration tested.

**Figure 5-S2:**
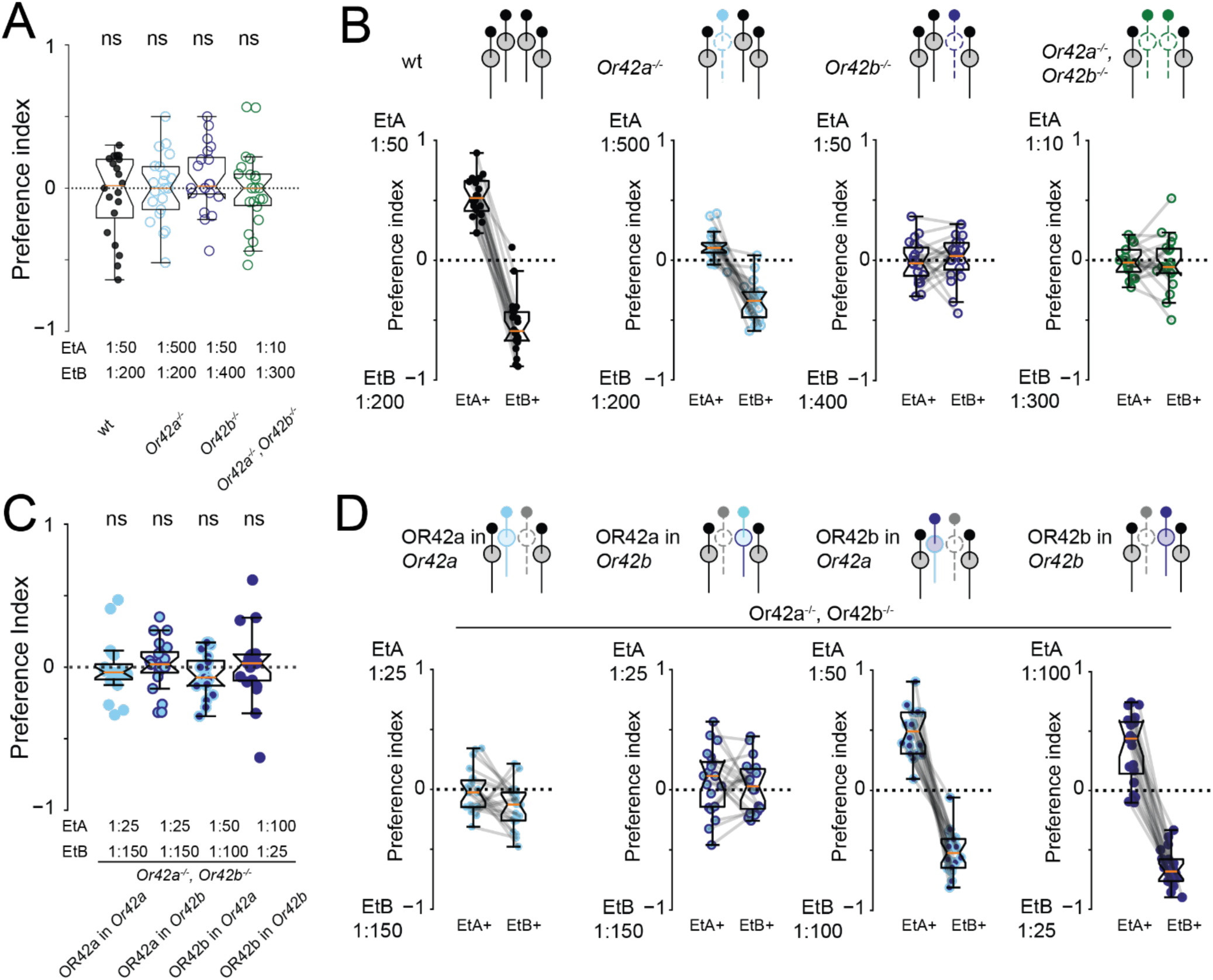
OR42b is necessary and sufficient for EtA vs. EtB discrimination. **(A)** Equal naïve preference in receptor knockout experiments (wt n=20, *Or42a*^-/-^ n=21, *Or42b*^-/-^ n=20, Del(*Or42a,Or42b*) n=22, one-sample *t* test followed by Holm-Bonferroni correction, ns p>0.05) **(B)** Reciprocal preference indices underlying discrimination results in Figure 5C. **(C)** Equal naïve preference in receptor rescue experiments (n=20, Wilcoxon signed-rank test with Holm-Bonferroni correction, ns p>0.05). **(D)** Reciprocal preference indices underlying discrimination results in Figure 6.

**Figure 6-S1:**
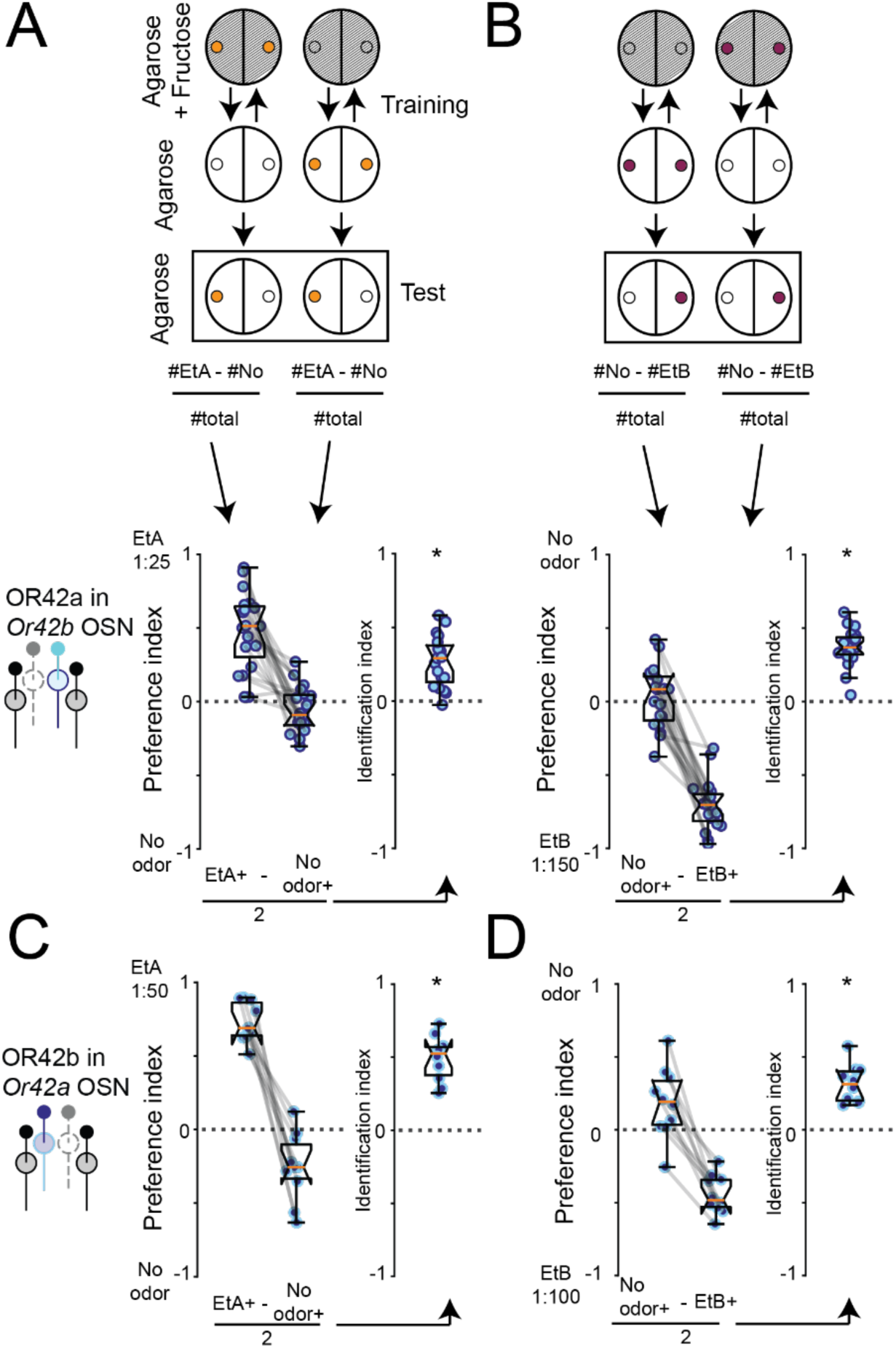
Larvae with ectopic receptor expression associate all tested odors with reward. **(A,B)** Larvae expressing OR42a in *Or42b* OSN associate presence or absence of EtA and EtB with reward (n=19, one-sample *t* test, *p<0.05). **(C, D)** Larvae expressing OR42b in *Or42a* OSN associate presence and absence of EtA and EtB with reward (n=10, one-sample *t* test, *p<0.05). These results confirm that discrimination deficits reflect OSN activity patterns rather than learning impairments (see Methods).

